# Branched chain α-ketoacids aerobically activate HIF1α signaling in vascular cells

**DOI:** 10.1101/2024.05.29.595538

**Authors:** Wusheng Xiao, Nishith Shrimali, William M. Oldham, Clary B. Clish, Huamei He, Samantha J. Wong, Bradley M. Wertheim, Elena Arons, Marcia C. Haigis, Jane A. Leopold, Joseph Loscalzo

**Author notes:** Corresponding author: Joseph Loscalzo, M.D., Ph.D. Department of Medicine Brigham and Women’s Hospital, 75 Francis St, Boston, MA 02115, USA.

## Abstract

Hypoxia-inducible factor 1α (HIF1α) is a master regulator of numerous biological processes under low oxygen tensions. Yet, the mechanisms and biological consequences of aerobic HIF1α activation by intrinsic factors, particularly in primary cells remain elusive. Here, we show that HIF1α signaling is activated in several human primary vascular cells under ambient oxygen tensions, and in vascular smooth muscle cells (VSMCs) of normal human lung tissue, which contributed to a relative resistance to further enhancement of glycolytic activity in hypoxia. Mechanistically, aerobic HIFα activation is mediated by paracrine secretion of three branched chain α-ketoacids (BCKAs), which suppress prolyl hydroxylase domain-containing protein 2 (PHD2) activity *via* direct inhibition and *via* lactate dehydrogenase A (LDHA)-mediated generation of L-2-hydroxyglutarate (L2HG). Metabolic dysfunction induced by BCKAs was observed in the lungs of rats with pulmonary arterial hypertension (PAH) and in pulmonary artery smooth muscle cells (PASMCs) from idiopathic PAH patients. BCKA supplementation stimulated glycolytic activity and promoted a phenotypic switch to the synthetic phenotype in PASMCs of normal and PAH subjects. In summary, we identify BCKAs as novel signaling metabolites that activate HIF1α signaling in normoxia and that the BCKA-HIF1α pathway modulates VSMC function and may be relevant to pulmonary vascular pathobiology.

## INTRODUCTION

Hypoxia-inducible factor 1α (HIF1α) is a master regulator of biological processes in hypoxia. Yet, the mechanisms and biological consequences of aerobic HIF1α activation by intrinsic factors, particularly in normal (primary) cells remain elusive. Here, we show that HIF1α signaling is activated in several human primary vascular cells in normoxia, and in vascular smooth muscle cells (VSMCs) of normal human lungs. Mechanistically, aerobic HIF1α activation is mediated by paracrine secretion of three branched chain α-ketoacids (BCKAs), which suppress PHD2 activity *via* direct inhibition and *via* LDHA-mediated generation of L-2-hydroxyglutarate. BCKA-mediated HIF1α signaling activation stimulated glycolytic activity and governed a phenotypic switch of pulmonary artery SMCs, which correlated with BCKA metabolic dysregulation and pathophenotypic changes in pulmonary arterial hypertension patients and male rat models. We thus identify BCKAs as novel signaling metabolites that aerobically activate HIF1α and that the BCKA-HIF1α pathway modulates VSMC function and may be relevant to pulmonary vascular pathobiology.

Hypoxia-inducible factor 1 (HIF1) is a central regulator of cellular adaptive and survival responses to a low oxygen environment through governing hundreds of gene transcripts involved in energy metabolism, angiogenesis, erythropoiesis, vascular tone, cell growth and survival, cell stemness, autophagy, redox homeostasis, embryonic development, and others^1,2^. HIF1 functions as a heterodimer that consists of an oxygen-sensitive alpha subunit (HIF1α) and an oxygen-insensitive beta subunit (HIF1β)^3,4^. Under ample oxygen conditions, HIF1α protein is first hydroxylated on two proline residues (Pro402 and Pro564) by prolyl hydroxylase domain-containing proteins (PHD1-3) and subsequently recognized by von-Hippel Lindau protein (pVHL) in an E3 ubiquitin ligase complex resulting in polyubiquitination and proteasomal degradation^4^. Thus, dysfunction of this mechanistic machinery, *e.g.,* inactivation of PHD enzymes, could lead to HIF1α protein stabilization and activation of its downstream targets to modulate numerous (patho)physiological conditions.

The PHD dioxygenases contain Fe^2+^ as an active center and require oxygen, α-ketoglutarate (α-KG), and ascorbate as substrates^3,4^. In addition to decreased oxygen levels, a well-documented mechanism, emerging evidence supports inhibition of PHD activity by limiting other required substrates including iron chelators (*e.g.,* desferrioxamine), oxidization of Fe^2+^ and ascorbate by reactive oxygen species (ROS), and competition with α-KG^4^. For example, the intermediary metabolites of the tricarboxylic acid (TCA) cycle succinate and fumarate have been shown to activate HIF1α signaling by competitively inhibiting the binding of α-KG to PHDs and by covalently binding with glutathione to increase mitochondrial ROS production^5–8^. Glycolytic end-products pyruvate and lactate may also stabilize HIF1α protein and enhance its transcriptional activity by an unknown mechanism(s) in aerobic cancer cells^9^. Recently, Burr and colleagues showed that aerobic accumulation of L-2-hydroxylglutarate (L2HG), but not its enantiomer D2HG, inhibits PHD hydroxylase activity leading to stabilization of HIF1α protein in human KBM-7 leukemia and HeLa cells when mitochondrial α-KG dehydrogenase (KGDH) complex is depleted^10^. Furthermore, in triple negative breast cancer cells, glutamate accumulation was able to inhibit PHD2 activity *via* a redox mechanism by blockade of extracellular cystine uptake and depletion of intracellular cysteine levels, consequently resulting in activation of HIF1α under normoxia^11^. Of interest, these known oxygen-independent activation mechanisms of HIF1α signaling were exclusively observed in aerobic cultures of cancer cells.

Yet, aerobic activation of HIF1α signaling and its underlying mechanisms remain poorly understood in normal (primary) cells. Given the facts that HIF1α is a crucial regulator of angiogenesis, vascular homeostasis and function, and vascular pathogenesis^12,13^, and that manipulating HIF1α activity serves as a promising therapeutic strategy for the treatment of cardiopulmonary vascular diseases^13,14^, we investigated the possible mechanisms of aerobic HIF1α activation in normal (primary) vascular smooth muscle cells (VSMCs) and its relevance to the pathobiology of pulmonary arterial hypertension (PAH), a disease characterized by HIF1α activation, increased mean pulmonary arterial pressure (mPAP) and pulmonary vascular resistance (PVR), and obliterative vascular remodeling^15^. We discovered that branched chain α-ketoacids (BCKAs), metabolites of branched chain amino acids (BCAAs; leucine, isoleucine, and valine), activate HIF1α signaling under aerobic conditions through iron chelation and L2HG-dependent inhibition of PHD2 activity, and that BCKA metabolism is dysregulated in PAH animal models and human patients, highlighting the BCKA-HIF1α pathway in the regulation of VSMC function and as a possible therapeutic target for select vascular diseases.

## Results

### Aerobic activation of HIF1α buffers hypoxia-induced glycolytic responses in VSMCs

To explore aerobic activation of HIF1α signaling in normal (*i.e.,* not immortalized) cells, we first examined HIF1α protein levels in aerobic cultures of commercially available primary human vascular cells and fibroblasts and compared them with human cancer cell lines. We found that HIF1α protein was detectable in normoxic cultures of human pulmonary arterial SMCs (PASMCs) and aortic SMCs (AoSMCs) but not endothelial cells of different vascular beds, lung fibroblasts, dermal fibroblasts, or cancer cell lines under our experimental conditions (**Fig. 1a**). This finding appears to be independent of *HIF1α* transcript abundance as its mRNA levels in PASMCs and endothelial cells were comparable (data not shown). Confocal microscopy further demonstrated that HIF1α protein was present in the nucleus of PASMCs but was absent in endothelial cells in normoxia (**Fig. 1b**), and that HIF1α protein was predominately detected in VSMCs of normal human lung tissue (**Fig. 1c**). Consistently, aerobic HIF1α stabilization in VSMCs correlated with relatively higher basal mRNA expression of its known target genes in glycolysis, including *glucose transporter 1* (*GLUT1*), *hexokinase 2* (*HK2*), *6-phosphofructo-2-kinase/fructose 2,6-biphosphatase 3* (*PFKFB3;* to a lesser extent), and *lactate dehydrogenase A* (*LDHA*), than the other seven cell types screened (Extended Data Fig. 1a-d). These gene expression profiles were consistent with the high basal glycolytic activity in PASMCs and AoSMCs, as their extracellular acidification rate (ECAR; indicative of glycolysis) was higher than that of three types of endothelial cells and dermal fibroblasts, despite it being lower than two cancer cell lines studied (likely a consequence of increased metabolic demand by these rapidly proliferating immortalized cell lines) (**Fig. 1d**). These results support that HIF1α signaling is constitutively active in VSMCs under aerobic conditions.

**Fig 1.**
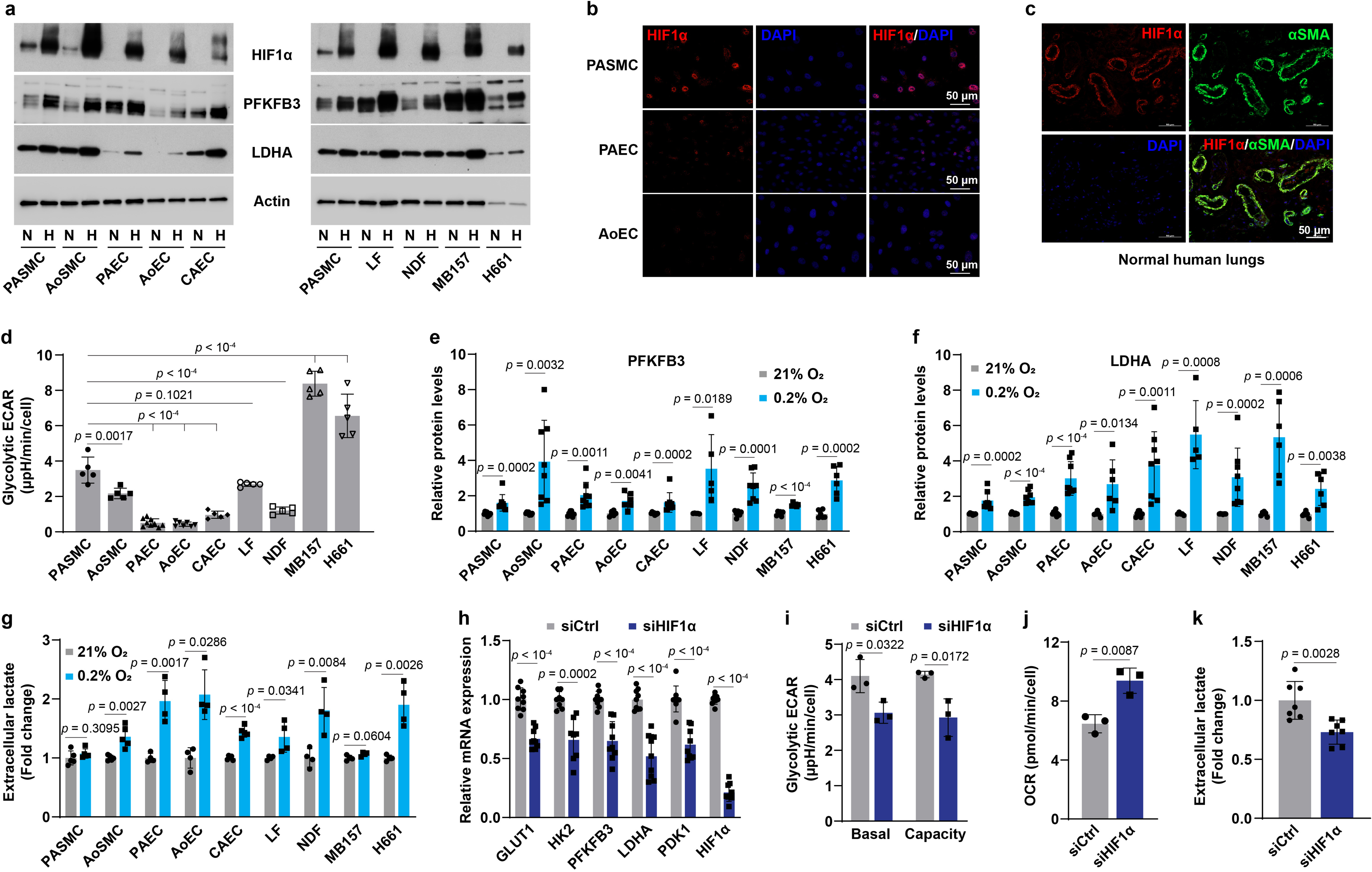
Aerobic HIF1α stabilization in VSMCs. **a,** Immunoblotting images showing the protein levels of HIF1α and its targets in 9 types of normal and cancer cells under normoxia (N) and hypoxia (H; 0.2% O_2_ for 24 hours). *n* = 5-8. **b,** Confocal microscopy shows HIF1α expression in aerobic cell cultures of human PASMCs, PAECs, and AoECs. *n* = 3. **c,** HIF1α and α-smooth muscle actin (αSMA, a marker of VSMCs) expression in human lung tissues of transplantation-failed donors. *n* = 3. **d,** Glycolytic extracellular acidification rate (ECAR) in normoxic cell cultures determined by a Seahorse glycolytic stress assay. *n* = 5-9. **e,f,** PFKFB3 (**e**) and LDHA (**f**) protein levels in cells cultured in 21% or 0.2% O_2_ for 24 hours. Fold change was calculated relative to corresponding normoxic cultures of each cell type. *n* = 5-8. **g,** Extracellular lactate levels in different normal and cancer cell types under normoxia or hypoxia (0.2% O_2_) for 24 hours. *n* = 4-5. **h,** mRNA expression of HIF1α and its known transcriptional genes in aerobic cultures of PASMCs transfected with control siRNA (siCtrl) or *HIF1α* siRNA (siHIF1α). *n* = 9. **i,j,** Seahorse assays show ECAR (**i**) and oxygen consumption rate (OCR; **j**) in PASMCs with *HIF1α* knockdown under normoxia. *n* = 3. **k,** Lactate secretion by PASMCs with *HIF1α* knockdown. Fold change was calculated relative to siCtrl-transfected cells. *n* = 7. All data were presented as mean ± SD. One-way ANOVA followed by Dunnett’s post-hoc test (**d**), Student’s t test or Mann-Whitney U test (**e-k**) was used when compared to aerobic culture of PASMCs or the matched cell type (**d-g**), or siCtrl-transfected PASMCs (**h-k**).

These findings led us to investigate the glycolytic response of these cells to hypoxia in more detail. Intriguingly, VSMCs, particularly PASMCs, showed the least upregulation of 4 glycolytic genes, GLUT1, HK2, PFKFB3, and LDHA, under hypoxia (**Fig. 1a, e, f**, and Extended Data Fig. 1e-h). Functionally, hypoxia induced much less of an increase in lactate secretion by PASMCs and AoSMCs when compared to other normal cell types and H661 lung cancer cells (**Fig. 1g**), consistent with the notion that high aerobic HIF1α activity contributes to the comparative resistance of VSMCs to hypoxia-induced enhancement of glycolytic activity.

These genetic and metabolic phenotypes relied on HIF1α activity since knockdown of *HIF1α* suppressed the transcription levels of its responsive genes in glucose metabolism, which correlated with decreases in basal glycolytic ECAR levels, glycolytic capacity, and lactate secretion, as well as a concomitant elevation in oxygen consumption rate (OCR, indicative of mitochondrial oxidative phosphorylation) (**Fig. 1h-k**). Overall, these observations indicate that HIF1α signaling is activated in VSMCs in normoxia, and that such high HIF1α basal activity impairs further hypoxia-induced glycolytic enhancement.

### Aerobic HIF1α stabilization is a paracrine phenomenon

Next, we sought to elucidate how HIF1α protein is stabilized under well-oxygenated conditions, and we hypothesized that aerobic HIF1α activation could be a (metabolic) paracrine effect in PASMCs. To test this hypothesis, conditioned medium (CM) was used to feed fresh PASMC subcultures (**Fig. 2a**). PASMCs cultured in CM expressed markedly higher levels of HIF1α protein and its transcriptional targets than cells cultured in regular growth medium (GM) (**Fig. 2b, c**). HIF1α protein accumulation significantly decreased after dilution of CM (Extended Data Fig. 2a), implying secreted molecules mediate HIF1α protein stabilization. Notably, CM specifically stabilized HIF1α but not HIF2α, which correlated with an over 3-fold higher basal *HIF1α* mRNA abundance than that of *HIF2α* in GM-cultured PASMCs (Extended Data Fig. 2b, c). In addition, such selective stabilization of HIF1α could be also explained by the fact that PHD2 is known to have different effects on HIF1α and HIF2α stability owing to HIF-associated factor facilitating stabilization of HIF1α but not HIF2α^16^ and of HIF1α induction of PHD3 which, in turn, blunts the stabilization of HIF2α^17^. Temporal experiments revealed that CM-mediated HIF1α protein stabilization peaked after 2-4 hours of incubation, while its target gene expression lagged, peaking after 8-14 hours (Extended Data Fig. 2d, f), also consistent with a paracrine phenomenon governing downstream gene transcription. Furthermore, HIF1α induction by CM seemed to be independent of its transcription and of proteins involved in HIF1α’s degradation, PHD2 and pVHL protein levels (Extended Data Fig. 2d, e, g). Rather, this effect is likely due to suppression of PHD2 enzymatic activity since the level of hydroxylated HIF1α was substantially lower in CM-cultured PASMCs than that in time-matched GM-cultured cells (Extended Data Fig. 2h).

**Fig 2.**
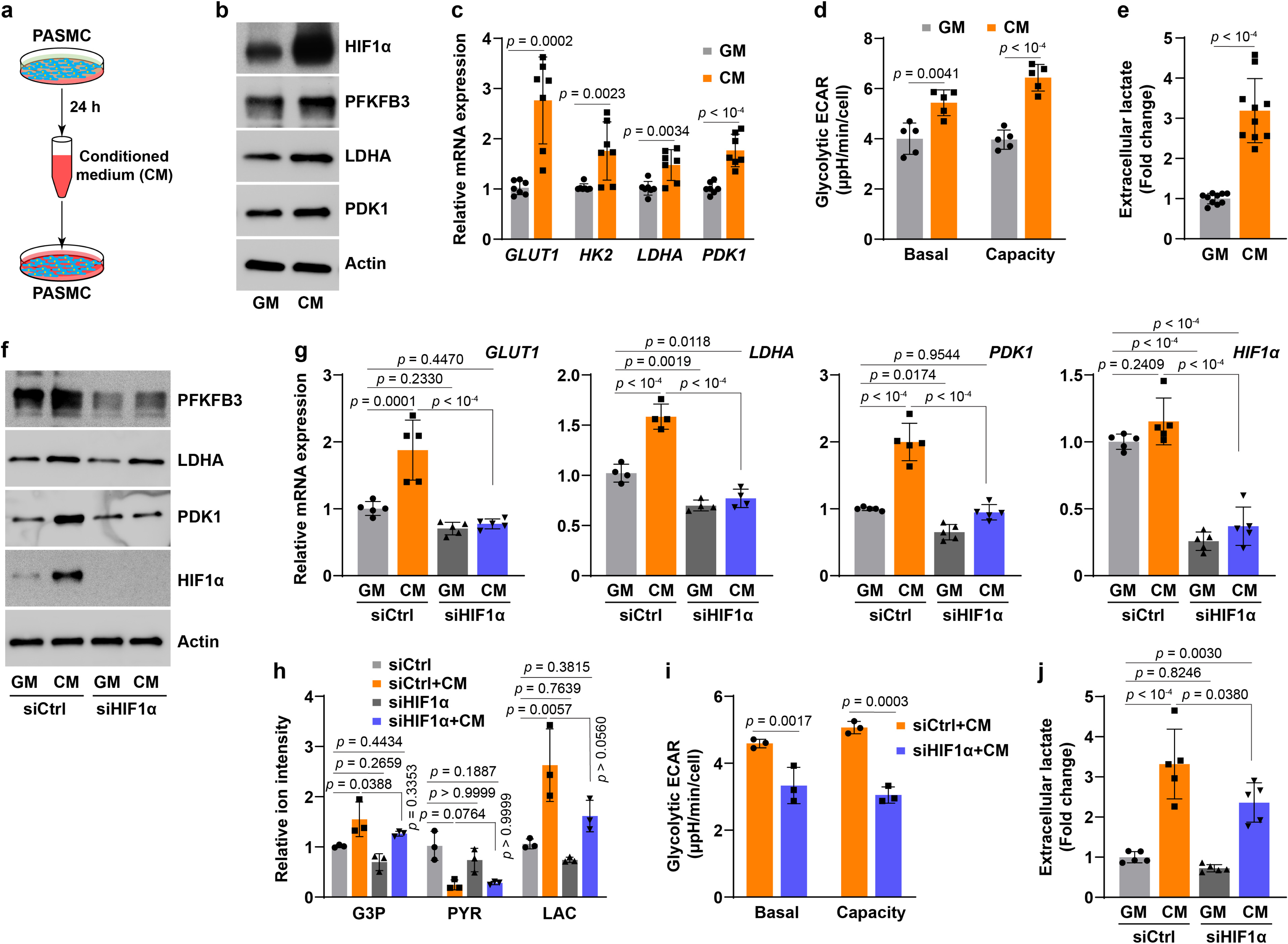
Aerobic activation HIF1α signaling in PASMCs is a paracrine effect. **a,** Schematics shows medium conditioning and reapplication. **b,** Representative immunoblots of HIF1α and its regulatory proteins in PASMCs cultured in fresh growth medium (GM) or conditioned medium (CM). *n* = 3. **c,** Relative mRNA expression of HIF1α key target genes in glucose metabolism in PASMCs cultured in GM or CM. *n* = 7. **d,** Seahorse glycolytic stress test shows basal ECAR and glycolytic capacity in GM- or CM- cultured PASMCs. *n* = 5. **e,** Extracellular lactate levels in PASMCs grown in GM or CM. Fold change was calculated relative to GM-cultured cells. *n* = 10. **f,g,** Protein levels (**f**) and mRNA expression (**g**) of HIF1α and its transcriptional targets in PASMCs grown in GM or CM after transfection with control siRNA (siCtrl) or human *HIF1α* siRNA (siHIF1α). *n* = 3 (**f**) and 5 (**g**). **h,** LC-MS metabolomic profiling shows the levels of glycolytic metabolites G3P, pyruvate (PYR), and lactate (LAC) in PASMCs treated as described in panel **f**. *n* = 3. **i,** Seahorse glycolytic stress test shows basal ECAR and glycolytic capacity in CM-cultured PASMCs transfected with siCtrl or siHIF1α. *n* = 3. **j,** Extracellular lactate levels in GM or CM cultures of PASMCs with or without siHIF1α transfection. *n* = 5. All data are presented as mean ± SD. Student’s t test or Mann-Whitney U test (**c-e**, **i**), one-way ANOVA followed by Tukey’s post-hoc or Kruskal-Wallis test followed by Dunn’s post-hoc test (**g**, **h**, **j**) was used when compared to GM-cultured PASMCs (**c-e**), GM-cultured and siCtrl-transfected PASMCs (**g**, **h**, **j**), or siCtrl-transfected and CM-cultured PASMCs (**g-j**).

We examined the metabolic consequences of CM-mediated HIF1α activation and found that glycolytic activity was enhanced in CM-cultured PASMCs, as evidenced by ∼1.5-fold increases in glycolytic ECAR and capacity, as well as a 3-fold increase in lactate secretion (**Fig. 2a-e**). Strikingly, these CM-mediated metabolic changes were dependent on HIF1α activity as silencing *HIF1α* significantly abrogated the upregulated mRNA and protein expression of HIF1α target genes (**Fig. 2f, g**). As a result, *HIF1α* knockdown mitigated the increases in glycolytic products glyceraldehyde 3-phosphate (G3P) and lactate, albeit the reduction of pyruvate levels remained (**Fig. 2h**). Moreover, the enhanced glycolytic activity and the elevated lactate secretion were significantly attenuated in CM-cultured and *HIF1α*-silenced PASMCs (**Fig. 2i, j**), supporting the conclusion that secreted molecule(s) in CM stimulate(s) glycolytic activity in PASMCs *via* a HIF1α-dependent mechanism. Therefore, aerobic HIF1α protein stabilization and activation in PASMCs may be mediated by a paracrine molecule(s).

### Branched chain α-ketoacids (BCKAs) mediate paracrine activation of HIF1α signaling

To identify the chemical nature of the potential paracrine mediator(s), we found that it was not inactivated by boiling or proteinase K treatment of CM, and that it was retained in filtrates of less than 10 kDa in size (Extended Data Fig. 2i-l). Thus, we speculated that the potential factor(s) is a small molecule. Based on previous reports^4,9^, we first screened and ruled out lactate, pyruvate, fumarate, succinate, aspartate, and malate as the aerobic stabilizers of HIF1α in PASMCs since no (dose-dependent) inductions of HIF1α and its target pyruvate dehydrogenase kinase 1 (PDK1) proteins were observed (Extended Data Fig. 2m-o).

We next performed LC-MS metabolomic profiling in CM derived from PASMCs to identify potential mediators, and found that α-ketoisocaproate (KIC), α-keto-β-methylvalerate (KMV), and α-ketoisovalerate (KIV) were the top 3 most increased metabolites (**Fig. 3a, b**). Consistently, a 2-3-fold increase in intracellular KIC, KMV, and KIV levels was also observed in CM-cultured PASMCs (**Fig. 3c**). These metabolites--KIC, KMV, and KIV--are the ketoacid derivatives (collectively denoted BCKAs) of the three BCAA leucine, isoleucine, and valine, respectively. Dysregulation of BCAA catabolism has been documented in many cardiovascular metabolic diseases^18,19^; however, little is known about the regulatory effects of BCKAs on HIF1α protein stability. Thus, we treated PASMCs with millimolar concentrations of each BCKAs (concentrations found in the plasma of maple syrup urine disease patients^20^). Strikingly, we observed notable stabilization of HIF1α protein and increases in the mRNA expression of *HK2*, *GLUT1*, and *PDK1* (Extended Data Fig. 3a, d). Importantly, such effects were not seen in cells treated with butyrate, the fourth most enriched metabolite in CM (**Fig. 3a** and Extended Data Fig. 3a, d). Neither BCKAs nor butyrate were able to induce HIF2α protein in PASMCs (Extended Data Fig. 3b), indicating that BCKAs specifically stabilize HIF1α protein.

**Fig 3.**
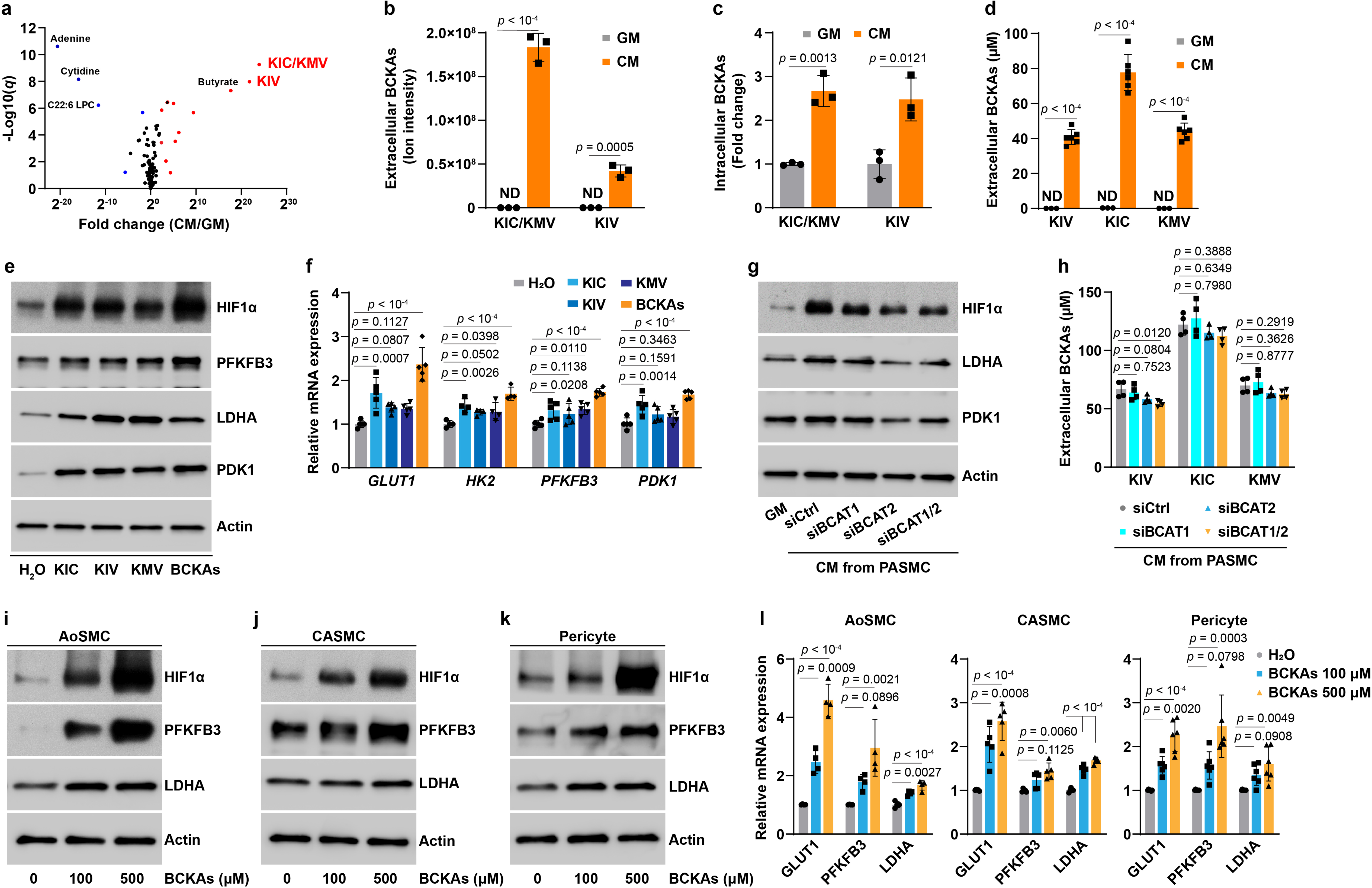
Secreted BCKAs in CM activate HIF1α signaling under aerobic conditions. **a,** Volcano plot of LC-MS metabolomic profiling shows the levels of 138 secreted metabolites in CM of PASMCs. *n* = 3. **b,** Quantitation of KIC/KMV (both have the same m/z ratio and are indistinguishable by LC-MS) and KIV in CM of PASMCs. ND: non-detectable. *n* = 3. **c,** Intracellular levels of KIC/KMV and KIV by LC-MS analysis in PASMCs cultured in GM or CM. Fold change was calculated relative to GM-cultured cells. *n* = 3. **d,** HPLC measurements of BCKAs in CM of PASMCs and basal GM. *n* = 3-6. **e,f,** Protein levels (**e**) and mRNA expression (**f**) of HIF1α and its target genes in PASMCs treated with KIC (100 μM), 50 μM of KMV or KIV, or all three BCKAs combined (KIC 100 μM, 50 μM KMV, and 50 μM KIV). *n* = 3 (**e**) and 5 (**f**). Fold change in **f** was calculated relative to vehicle control H_2_O treated cells. **g,** Protein levels of HIF1α, LDHA, and PDK1 in PASMCs cultured in GM or CM from PASMCs transfected with *siRNAs* for control (siCtrl), *BCAT1* (siBCAT1), *BCAT2* (siBCAT2), or both *BCAT1* and *BCAT2* (siBCAT1/2). *n* = 3. **h,** HPLC determination of secreted BCKA levels in CM of PASMCs after transfection with siCtrl, siBCAT1, siBCAT2, or siBCAT1/2. *n* = 4. **i-k,** Representative immunoblotting images showing HIF1α and its target proteins PFKFB3 and LDHA in BCKA-treated human AoSMCs (**i**), CASMCs (**j**), and pericytes (**k**). *n* = 3. **l,** mRNA expression of HIF1α transcriptional genes in glycolysis in BCKA-treated human AoSMCs (*n* = 4), CASMCs (*n* = 5), and pericytes (*n* = 6). Fold change was calculated relative to untreated control cells. All data are presented as mean ± SD. Student’s t test (**b-d**), one-way ANOVA followed by Dunnett’s test or Kruskal-Wallis test followed by Dunn’s test (**f**, **h**, l) was applied when compared to GM (**b-d**), vehicle control H_2_O treated (**f**, **l**), or CM from siCtrl-transfected PASMCs (**h**).

Since the concentrations of BCKAs in CM were the key factors to determine whether these metabolites were the authentic mediators of aerobic HIF1α stabilization, using HPLC measurements we detected roughly 80 μM KIC and 40 μM for KIV and KMV in CM (**Fig. 3d**), and thus treated PASMCs with comparable doses of BCKAs (100 μM of KIC, 50 μM of either KIV or KMV, or all 3 combined). In keeping with the results using millimolar doses (Extended Data Fig. 3a, d), lower doses of BCKAs were sufficient to upregulate HIF1α protein and its responsive gene expression (**Fig. 3e, f** and Extended Data Fig. 3c, d). Temporal experiments showed that BCKA-induced HIF1α stabilization and upregulation of its downstream targets were observed as early as 6 hours and continued up to 24 hours; by contrast, no induction was noted in HIF2α protein (Extended Data Fig. 3e-g). These results led us to conclude that BCKAs are the principal mediators of CM-induced activation of HIF1α signaling in PASMCs.

This conclusion was further reinforced by using CM collected from cells in which the two branched chain amino acid transaminases (cytosolic *BCAT1* and mitochondrial *BCAT2*) were knocked down, both of which catalyze the reversible conversion of BCAAs to BCKAs^18^.

Interestingly, we found that the CM-mediated activation of HIF1α signaling activity was significantly dampened in PASMCs cultured in CM from *BCAT2* alone or *BCAT1/2* double silenced PASMCs, whereas no effects were observed in CM from *BCAT1* knockdown cells (**Fig. 3g** and Extended Data Fig. 3h, i). This interpretation was consistent with the fact that the secreted levels of KIV (to a greater extent), KIC, and KMV in CM were lower in PASMCs with *BCAT2* alone or *BCAT1/2* double but not *BCAT1* silencing compared to that of control cells (**Fig. 3h**), indicating BCAT2-catalyzed production of BCKAs is the dominant contributor to CM-induced HIF1α activation. Furthermore, aerobic HIF1α protein stabilization vanished in PASMCs cultured in BCAA-free medium, while exogenous supplementation of BCKAs restored HIF1α protein stabilization and its transcriptional activity (Extended Data Fig. 3j, k), supporting the requirement of BCKAs for aerobic HIF1α signaling activity. In summary, we identify that BCKAs, particularly KIV, produced by BCAT2 are the paracrine mediators of aerobic HIF1α activation.

To examine whether induction of aerobic stabilization of HIF1α by BCKAs is true for other cell types, we screened 12 primary cells and 7 cancer cell lines. Surprisingly, BCKA treatment dose-dependently stabilized HIF1α protein and induced the expression of PFKFB3, LDHA, and GLUT1 in human AoSMCs, coronary artery SMCs (CASMCs), placental pericytes, and colorectal adenocarcinoma Caco2 cells (a relatively modest effect) (**Fig. 3i-l** and Extended Data Fig. 3l, m); however, the same treatment regimen failed to do so in the other 15 cell types tested under our experimental conditions (Extended Data Fig. 3n and data not shown). One common feature that was shared by these responsive cell types and distinct from those non-responsive cell types was the readily detectable HIF1α protein in aerobic control cells (**Fig. 3e, i-k** and Extended Data Fig. 3l). Such discrepancies seemed to be independent of the levels of phosphorylated branched chain ketoacid dehydrogenase protein (BCKDH), the rate-limiting enzyme irreversibly catalyzing oxidative decarboxylation of BCKAs (Extended Data Fig. 3o). Thus, these findings further support that BCKAs mediate the paracrine aerobic activation of HIF1α signaling in different cell types, but not as a uniformly generalizable phenomenon.

### BCKAs inhibit PHD2 hydroxylase activity through iron chelation and L2HG production

We next sought to ascertain the mechanism(s) responsible for BCKA-mediated aerobic HIF1α activation. By monitoring the levels of hydroxylated HIF1α protein after proteasome inhibition with MG132, we noted a delayed recovery of HIF1α hydroxylation in PASMCs treated with BCKAs, implying reduced PHD activity in these cells (Extended Data Fig. 4a). Consistently, results from a competitive inhibition assay showed that KIV dramatically inhibited PHD2 enzymatic activity, while KIC and KMV had more modest effects when used alone (**Fig. 4a**). PHD2 inhibition became more potent when the three BCKAs were used in combination, with an apparent IC_50_ of 252 μM, although their potency, while consistent with available intracellular concentrations, was lower than roxadustat (a known potent and selective PHD inhibitor with IC_50_ = 17 μM) (**Fig. 4a, b** and Extended Data Fig. 4b). Similar to α-KG and roxadustat, the predicted binding sites of BCKAs on PHD2 were most likely at the catalytic Fe^2+^ active center and perhaps at Tyr303 with estimated docking scores of -5.5∼-7.0 kcal/mol (**Fig. 4c** and Extended Data Fig. 4c), indicating that BCKAs are likely iron chelators that thereby inhibit PHD2 activity and stabilize HIF1α in normoxia.

**Fig 4.**
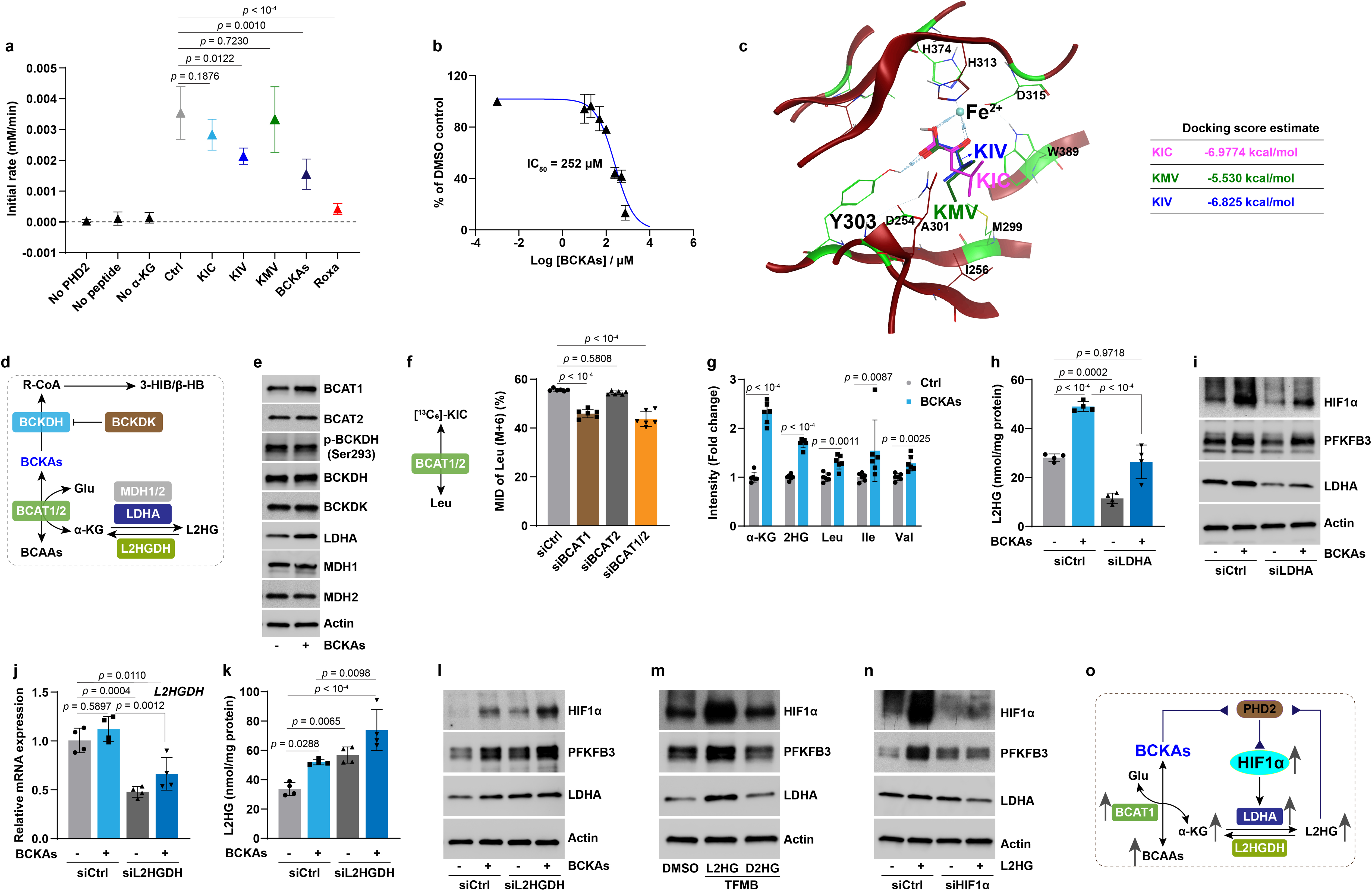
BCKAs inhibit PHD2 activity *via* a dual mechanism under normoxia. **a,** Reaction rates of KIC (250 μM), KMV (250 μM), KIV (250 μM), and all three BCKAs (200 μM of each BCKAs) determined by a competitive inhibition assay. Roxadustat (Roxa; 100 μM) was included as a positive control. *n* = 4. **b,** Inhibition curve and IC_50_ value of three BCKAs for PHD2 hydroxylase activity. *n* = 4. **c,** The predicted docking sites and energy of KIC, KMV, and KIV on PHD2 enzyme (crystal structure accession ID: 2G19). **d,** Schematic diagram shows the metabolic fates of BCKAs. **e,** Representative immunoblots in untreated and BCKA-treated PASMCs. *n* = 3. **f,** ^13^C_6_-KIC tracing of the labelled leucine (Leu) in PASMCs with *BCAT1*, *BCAT2*, or both (*BCAT1/2*) silencing. *n* = 6. **g,** LC-MS measurements of metabolite levels in BCKA-treated PASMCs. Fold change was calculated relative to untreated control cells. *n* = 6. **h,i,** Intracellular L2HG levels (**h**) and protein levels of HIF1α, PFKFB3, and LDHA (**i**) in siCtrl or *LDHA* siRNA (siLDHA) transfected PASMCs with or without BCKA treatment. *n* = 4 (**h**) and 3 (**i**). **j,** *L2HGDH* mRNA expression in siCtrl or *L2HGDH* siRNA (siL2HGDH) transfected PASMCs with or without BCKA treatment. *n* = 4. **k,l,** Intracellular L2HG levels (**k**) and protein levels of HIF1α, PFKFB3, and LDHA (**l**) in siCtrl or siL2HGDH transfected PASMCs with or without BCKAs. *n* = 4 (**k**) and 3 (**l**). **m,** Immunoblotting shows protein levels in cell permeable trifluoromethylbenzyl (TFMB) ester of L2HG (500 μM) or D2HG (500 μM) treated PASMCs. *n* = 3. **n,** Immunoblots of HIF1α, PFKFB3, and LDHA in siCtrl or *HIF1α* siRNA (siHIF1α) transfected PASMCs with or without cell permeable L2HG (TFMB-L2HG) (500 μM). *n* = 3. **o,** Proposed mechanisms of BCKA-induced inhibition of PHD2 activity. All data are presented as mean ± SD. One-way ANOVA followed by Dunnett’s test (**a**, **f**) or Tukey’s test (**h**, **j**, **k**), Student’s t test or Mann-Whitney U test (**g**) was used when compared to no inhibitor control (**a**), untreated control cells (**g**), siCtrl-transfected and control or BCKA-treated PASMCs (**f**, **h**, **j**, **k**).

Since the IC_50_ value of these BCKAs was much higher than the doses (50-100 μM) we used to stabilize HIF1α *in vitro*, we reasoned that a second inhibitory mechanism may exist and searched for it by examining the metabolic fates of BCKAs. Intracellular BCKAs can be irreversibly decarboxylated into R-CoA intermediates by the BCKDH complex and then into 3-hydroxyisobutyrate (3-HIB; KIV metabolite) and β-hydroxybutyrate (β-HB; KIC metabolite) (**Fig. 4d**). We first ruled out this possibility because the protein levels of BCKDH, its phosphorylated form (p-BCKDH), and its kinase (branched chain ketoacid dehydrogenase kinase, BCKDK) remained unchanged in BCKA-treated PASMCs (**Fig. 4e**), and because supplementation with 3-HIB or β-HB failed to stabilize HIF1α *in vitro* (data not shown).

We then focused on the second metabolic route whereby BCKAs are reversibly reaminated to BCAAs by BCAT1/2 in conjunction with the production of α-KG from glutamate (**Fig. 4d**). We and others previously reported that α-KG could be reduced to L2HG by spurious activity of LDHA and malate dehydrogenases (MDH1-2) in hypoxia^21,22^. L2HG is known as an inhibitor of PHD enzymes, stabilizing HIF1α^10,23–25^. However, whether this metabolic pathway is important under aerobic conditions remains unclear. We observed that BCAT1 and LDHA protein levels were selectively induced by BCKAs; by contrast, the protein levels of BCAT2, MDH1, and MDH2 were unaltered in PASMCs (**Fig. 4e**). In line with a recent report in cardiomyocytes^26^, using ^13^C_6_-KIC tracing, we found that roughly 60% of labeled KIC was reaminated to leucine, and that this reaction was mainly catalyzed by BCAT1 since silencing *BCAT1* alone or *BCAT1/2* combined was both equally efficient in reducing the proportion of leucine labeling, while knock down of *BCAT2* alone had no effect (**Fig. 4f**). As a result, BCKAs were able to increase the levels of all three BCAAs, α-KG, and total 2-hydroxyglutarate (2HG) significantly in PASMCs (**Fig. 4g**).

Since BCKAs, particularly KMV (at 10-20 mM dose range), are known inhibitors of KGDH resulting in elevation of intracellular α-KG levels^22,27,28^, the increase in α-KG level we observed led us to test the impact of BCKAs at physiological levels on KGDH activity. Interestingly, unlike the inhibition of KGDH by KMV at supraphysiological levels, an increasing trend of KGDH activity was seen in PASMCs treated with BCKAs at micromolar levels (Extended Data Fig. 4d), ruling out the possibility that the increased α-KG level in BCKA-stimulated cells was due to KGDH inhibition. For this reason, we next further focused on the elevated levels of total 2HG, derivatives of α-KG. The two enantiomers of 2HG (L2HG *vs.* D2HG) had opposite effects on PHD activity with L2HG being an inhibitor and D2HG being an agonist^25^. We, therefore, developed a colorimetric assay to measure L2HG levels specifically and found a 1.8-fold increase in L2HG levels by BCKA treatment, which was closely correlated with the 1.7-fold increase in total 2HG (**Fig. 4g, h**), in support of the idea that L2HG but not D2HG is specifically elevated by BCKAs. In short, these results support the conclusions that BCAT1-mediated reamination of BCKAs to BCAAs leads to accumulation of L2HG, which serves as a potent inhibitor of PHD2 to activate HIF1α signaling under normoxic conditions.

We next probed this model by modulating intracellular and extracellular L2HG levels with genetic and pharmacological approaches. The upregulation of LDHA by BCKAs in **Figure 4e** led us to hypothesize that LDHA is the central enzyme for L2HG generation. Indeed, silencing *LDHA* significantly reduced L2HG levels in both untreated and BCKA-treated PASMCs, which correlated with mitigation of BCKA-induced upregulation of HIF1α and PFKFB3 protein levels (**Fig. 4h, i**). By contrast, silencing *L2HG dehydrogenase* (L2HGDH), the solely known enzyme that metabolizes L2HG to α-KG^22^, was sufficient to increase intracellular L2HG levels in control cells and potentiate L2HG accumulation in BCKA-treated cells (**Fig. 4j, k**). Of note, the accumulation of L2HG closely correlated with the augmented activation of HIF1α signaling in control and BCKA-stimulated PASMCs (**Fig. 4l**). Furthermore, exogenous addition of cell permeable L2HG, but not D2HG, markedly induced HIF1α protein stabilization and its targets, PFKFB3 and LDHA expression (**Fig. 4m**). *HIF1α* silencing further validated that L2HG-induced PFKFB3 and LDHA upregulation depended on HIF1α activity (**Fig. 4n**). Thus, these data suggest that BCKAs inhibit PHD2 activity through direct iron chelation and LDHA-mediated L2HG generation resulting in aerobic activation of HIF1α signaling (**Fig. 4o**).

### BCKA-mediated aerobic HIF1α activation stimulates glycolytic activity

Metabolic reprogramming to glycolysis is a fundamental adaptive response upon HIF1α activation under low oxygen tensions^4^. We, thus, investigated the metabolic consequences of BCKA-mediated aerobic HIF1α activation. Metabolomic profiling and metabolite set enrichment analysis (MSEA) revealed that the significantly altered metabolites by BCKAs were mainly found in central glucose metabolism, including glycolysis, gluconeogenesis, the TCA cycle, and the Warburg effect (**Fig. 5a, b**). Specifically, the levels of glycolytic intermediates glucose 6-phosphate (G6P), dihydroxyacetone phosphate (DHAP), phosphoenolpyruvate (PEP), and pyruvate were significantly elevated upon BCKA incubation, while the increases of intracellular and extracellular lactate levels were modest (**Fig. 5c, d**). These metabolic changes correlated with upregulation of key glycolytic enzymes GLUT1, HK2, PFKFB3, and LDHA (**Fig. 5e, f**). It is notable that BCKA treatment also increased the TCA cycle intermediary metabolites with no effects on cellular ATP levels (**Fig. 4g** and Extended Data Fig. 5a, b), indicating that mitochondrial respiration remains functionally intact and active. This phenomenon was unique and differed from the dogma that a metabolic switch from mitochondrial respiration to glycolysis is induced by HIF1α under hypoxic conditions.

**Fig 5.**
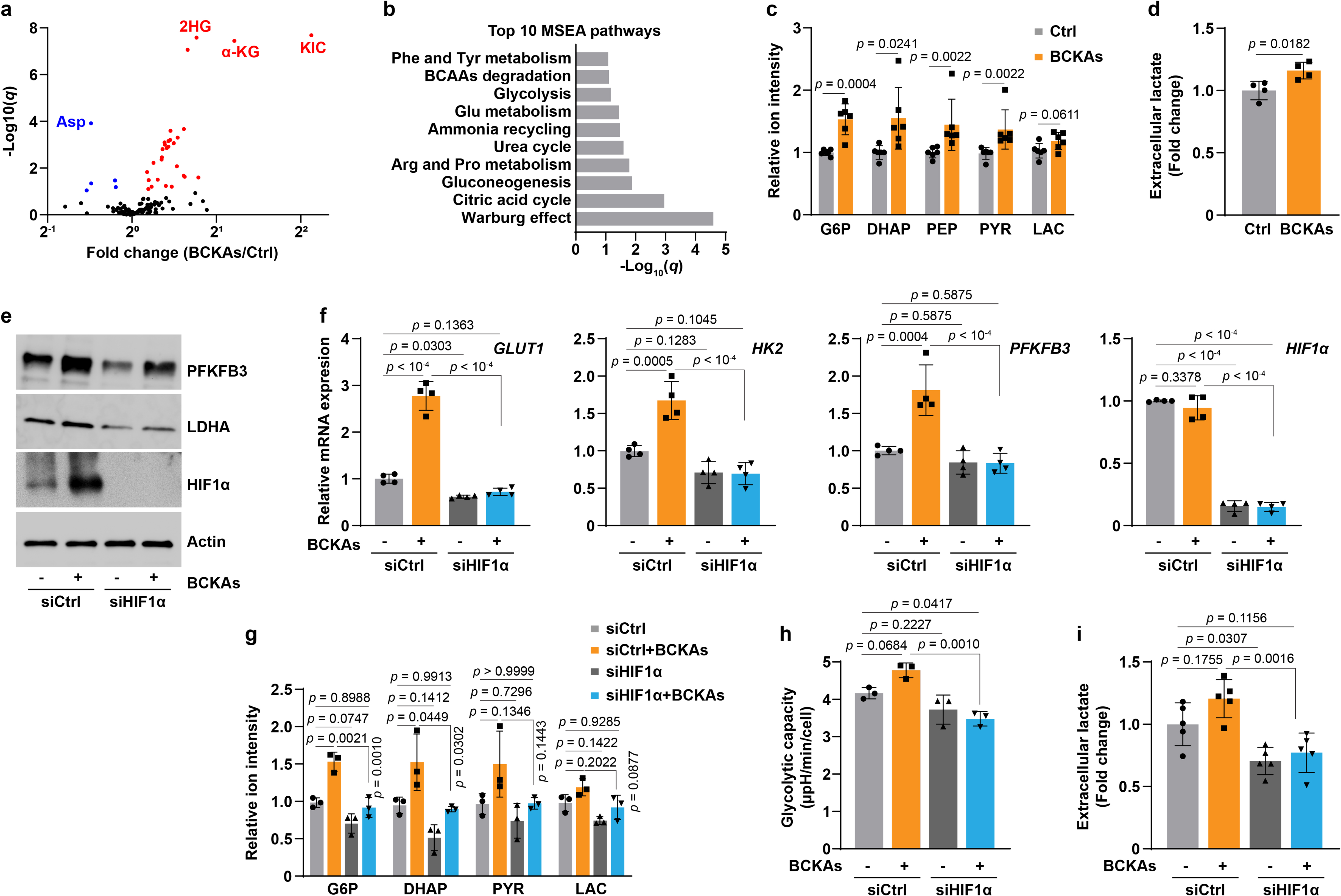
BCKAs enhance glycolysis *via* HIF1α activation. **a,** Volcano plot from LC-MS metabolomic profiling showing the changes of 136 metabolites in PASMCs with or without BCKA treatment. *n* = 6. **b,** Top 10 Metabolite Set Enrichment Analysis (MSEA) pathways in BCKA-treated PASMCs. **c,** LC-MS measurements of glycolytic metabolites in BCKA-treated PASMCs. Fold change was calculated relative to untreated controls. *n* = 6. **d,** Extracellular lactate levels in BCKA-treated PASMCs. Fold change was calculated relative to untreated controls. *n* = 4. **e,f,** Protein levels (**e**) and mRNA expression (**f**) of HIF1α and its transcriptional targets in PASMCs transfected with siCtrl or siHIF1α followed by stimulation with BCKAs. Fold change in **f** was calculated relative to control cells. *n* = 3 (**e**) and 4 (**f**). **g,** LC-MS measurements show the levels of glycolytic metabolites in PASMCs treated as described in panel **e**. Fold change was calculated relative to siCtrl-transfected and untreated controls. *n* = 3. **h,** Seahorse glycolytic stress test demonstrates glycolytic capacity of PASMCs treated as described in panel **e**. *n* = 3. **i,** Extracellular lactate levels in PASMCs treated as described in panel **e**. Fold change was calculated relative to siCtrl-transfected and untreated controls. *n* = 5. All data are presented as mean ± SD. Student’s t test or Mann-Whitney U test (**c**, **d**), or one-way ANOVA followed by Tukey’s test (**f-i**) was conducted when compared to untreated controls (**c**, **d**), siCtrl-transfected and untreated or BCKA-treated PASMCs (**f-i**).

Nevertheless, we next asked whether BCKA-mediated metabolic and genetic changes depended on HIF1α activity. As anticipated and consistent with the results in **Figure 2**, *HIF1α* knockdown abolished the induction of glycolytic genes by BCKAs (**Fig. 5e, f**). Consequently, the elevated levels of glycolytic metabolites, glycolytic capacity, and lactate secretion were also abrogated (to differing extents) in PASMCs with BCKA treatment and *HIF1α* silencing (**Fig. 5g-i**). By contrast, BCKA-induced changes in the TCA cycle metabolites and ATP derivatives [except aconitate (ACO), citrate (CIT), and ATP] were not dramatically affected by *HIF1α* silencing (Extended Data Fig. 5c, d). Thus, similar to hypoxia-induced HIF1α activation, aerobic activation of HIF1α signaling by BCKAs also enhances glycolytic activity and does so in a HIF1α-dependent manner.

### BCKA-mediated aerobic HIF1α activation regulates a phenotypic switch in VSMCs

VSMCs are essential for maintaining key physiological functions of the vasculature. Dysfunction of these cells, for example, phenotypic switching from contractile to synthetic subtype, have been implicated in the development of vascular diseases, such as atherosclerosis and PAH^29,30^. This phenotypic modulation is a fine-tuned process and governed by a repertoire of transcriptional regulatory mechanisms in response to internal and external stimuli, including oxygen tension^29,31^. For example, hypoxia (1% O_2_) promoted a synthetic phenotype (indicated by increased cell proliferation and migration and elevated synthetic marker gene expression) in rat PASMCs *via* HIF1α-mediated upregulation of microRNA-9^32^. Yet, no evidence is available regarding whether BCKAs could regulate VSMC phenotypes and whether such effects depend on HIF1α activation. Intriguingly, we found that BCKAs were able to stimulate expression of the synthetic marker gene *vimentin* (*VIM*) as effectively as that of platelet-derived growth factor (PDGF-BB; a widely used factor to promote synthetic phenotype) stimulation in human PASMCs, despite these agents having no effects on the expression of the contractile marker gene *α-smooth muscle actin* (*ACTA2*) as did transforming growth factor β (TGFβ; a known contractile phenotype stimulus) (**Fig. 6a**).

**Fig 6.**
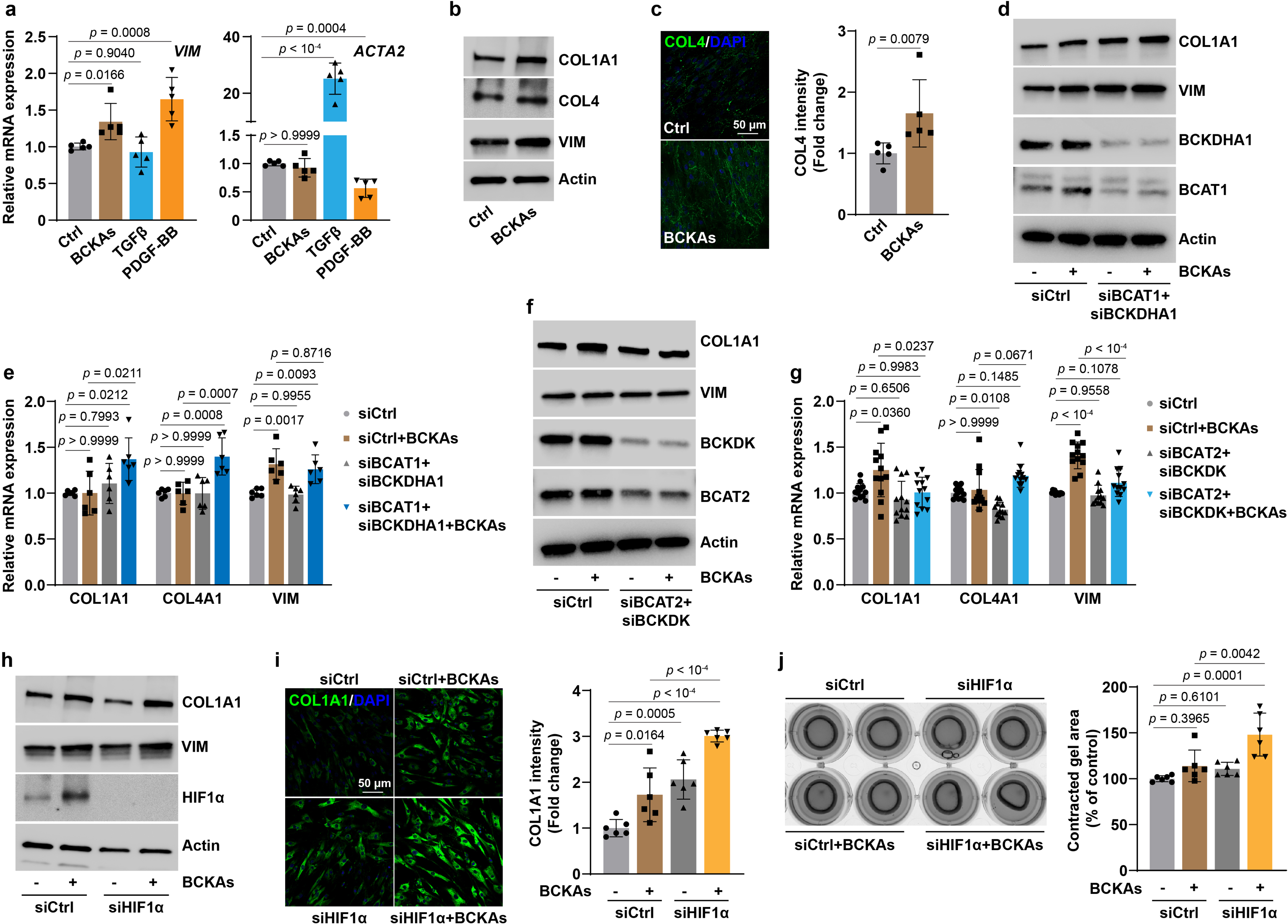
BCKAs promote a synthetic phenotype switch in PASMCs. **a,** mRNA expression of synthetic marker *VIM* and contractile marker *ACTA2* in BCKAs (100 μM), TGFβ (2 ng/mL), or PDGF-BB (10 ng/mL) stimulated cells. Fold change was relative to untreated controls. *n* = 5. **b,** Representative immunoblots of synthetic marker proteins in BCKA-stimulated cells. *n* = 6. **c,** Confocal microscopy images and quantitation represent COL4 synthesis and deposition in BCKA-treated PASMCs. Fold change was relative to control cells. *n* = 5. **d,e,** Protein (**d**) and mRNA (**e**) expression of synthetic marker genes in PASMCs transfected with control siRNA (siCtrl) or human *BCAT1* and *BCKDHA1* siRNAs (siBCAT1+siBCKDHA1) with or without BCKA treatment. Fold change was calculated relative to siCtrl-transfected and untreated cells. *n* = 3 (**d**) and 6 (**e**). **f,g,** Protein (**f**) and mRNA (**g**) expression of synthetic marker genes in PASMCs transfected with siCtrl or human *BCAT2* and *BCKDK* siRNAs (siBCAT2+siBCKDK) with or without BCKA treatment. Fold change was calculated relative to siCtrl-transfected and untreated cells. *n* = 3 (**f**) and 12 (**g**). **h,** COL1A1 and VIM protein expression in control and *HIF1*α knockdown PASMCs in the presence or absence of BCKAs. *n* = 8. **i,** Confocal microscopy and quantitation showing COL1A1 synthesis and deposition in control and *HIF1α* knockdown PASMCs in the presence or absence of BCKAs. Fold change was relative to control cells. *n* = 6. **j,** Collagen gel contraction assay illustrating the relative contracted area of collagen gels after 8-hour of detachment in control and *HIF1α* knockdown PASMCs with or without BCKAs. *n* = 6. All data are presented as mean ± SD. One-way ANOVA followed by Dunnett’s (**a**) or Tukey’s post-hoc test (**e**, **g**, **i**, **j**), or Kruskal-Wallis test followed by Dunn’s test (COL4A1 in **g**), or Mann-Whitney U test (**c**) was applied when compared to untreated controls (**a**, **c**), siCtrl-transfected and untreated or BCKA-treated PASMCs (**e**, **g**, **i**, **j**).

Furthermore, BCKAs also induced VIM, collagen 1A1 (COL1A1), and collagen 4 (COL4) protein expression and collagen biosynthesis by 1.3-2-fold (**Fig. 6b, c** and Extended Data Fig. 6a), supporting the conclusion that BCKAs promote a phenotypic switch toward synthetic subtype in human PASMCs.

To strengthen this viewpoint further, we manipulated intracellular BCKA levels by silencing their metabolic regulatory enzymes in conjunction with exogenous supplementation of these metabolites. As shown in **Figure 4**, BCAT1 catalyzes the reamination of BCKAs into BCAAs and the BCKDH complex catalyzes the irreversible decarboxylation of BCKAs. We, thus, knocked down both *BCAT1* and BCKDH catalytic E1 subunit α (*BCKDHA1*) to increase intracellular BCKA levels, and found that COL1A1 and VIM proteins were induced and that exogenous BCKA-induced upregulation of synthetic marker genes was heightened in PASMCs with the double knockdown of *BCAT1* and *BCKDHA1,* despite the fact that knockdown of these two genes *per se* had no effects on their mRNA expression levels (**Fig. 6d, e** and Extended Data Fig. 6b). By contrast, we also attempted to lower intracellular BCKA levels by knockdown of *BCAT2* and *BCKDK* since BCAT2 is the principal enzyme required for endogenous BCKA production (**Fig. 3**) and BCKDK blocks the downstream decarboxylation of BCKAs by phosphorylation and inhibition of the BCKDH complex. Notably, we found that silencing *BCAT2* and *BCKDK* abrogated BCKA-induced upregulation of COL1A1 and VIM (predominantly at its transcription level) expression, but had no effect on *COL4A1* transcription (**Fig. 6f, g** and Extended Data Fig. 6c). These findings, thus, indicate that BCKAs are mediators that promote and sustain the synthetic phenotype of PASMCs.

We next examined the involvement of HIF1α in these effects. Surprisingly, we found that knockdown of *HIF1α* further enhanced the upregulation of COL1A1 and VIM proteins and the increase in collagen biosynthesis, which led to greater contraction in these cells than in control cells (**Fig. 6h-j** and Extended Data Fig. 6d). Taken together, these data show that BCKAs promote a synthetic phenotype in PASMCs in which HIF1α activation appears to serve as a negative feedback mechanism to control collagen biosynthesis. Such an effect was further supported by the results showing that hypoxia (0.2% O_2_) dramatically activated HIF1α but repressed COL1A1 and COL4 expression in PASMCs (Extended Data Fig. 6e, f).

### Metabolic dysfunction of BCKAs in PAH pathophenotypes

These findings of BCKA-induced metabolic and phenotypic changes led us to investigate their potential relevance in PAH, a severe cardiopulmonary vascular disease characterized by HIF1α activation, enhanced glycolysis, increased collagen deposition and fibrosis, and PASMC dysfunction^15,30,33^. Correlation analyses from a PAH patient cohort (*n* = 32) revealed moderate positive correlation of the BCAA/BCKA ratio with mPAP (r = 0.32, *p* = 0.08) and with PVR (r = 0.35, *p* = 0.05), supporting a possible link between BCKA metabolism and PAH pathologies.

These findings led us to examine further such link in idiopathic PAH (IPAH) patient lungs. We found upregulation of key BCKA metabolic genes *BCAT1* (to a lesser extent), *BCAT2*, *BCKDK*, and *BCKDHA1* mRNA in the lungs of IPAH patients, which were accompanied with reduction of BCAT1 protein levels, and increases in BCAT2, BCKDK, and its substrate phosphorylated BCKDH (p-BCKDH) protein levels (**Fig. 7a, b** and Extended Data Fig. 7a). As the results in **Figures 3h** and **4f** suggested BCAT2 as the principal enzyme required for BCKA production and BCAT1 is the primary enzyme for BCAA synthesis, and since BCKDK-mediated inactivation of BCKDH blocks downstream catabolic steps of BCKAs, upregulation of BCAT2, BCKDK, and p-BCKDH proteins and downregulation of BCAT1 protein are expected to increase BCKA accumulation and decrease BCAA production in IPAH lungs. Indeed, KIC/KMV levels were significantly elevated in IPAH lung tissue with concomitant decreases in leucine and isoleucine (to a lesser extent) levels but an increase in valine levels (**Fig. 7c**), which correlated with a 2-3-fold elevation in HIF1α protein levels, predominantly in the VSMC layer (**Fig. 7d**), supporting a positive correlation between elevation of KIC/KMV levels and upregulation of HIF1α protein in IPAH lungs.

**Fig 7.**
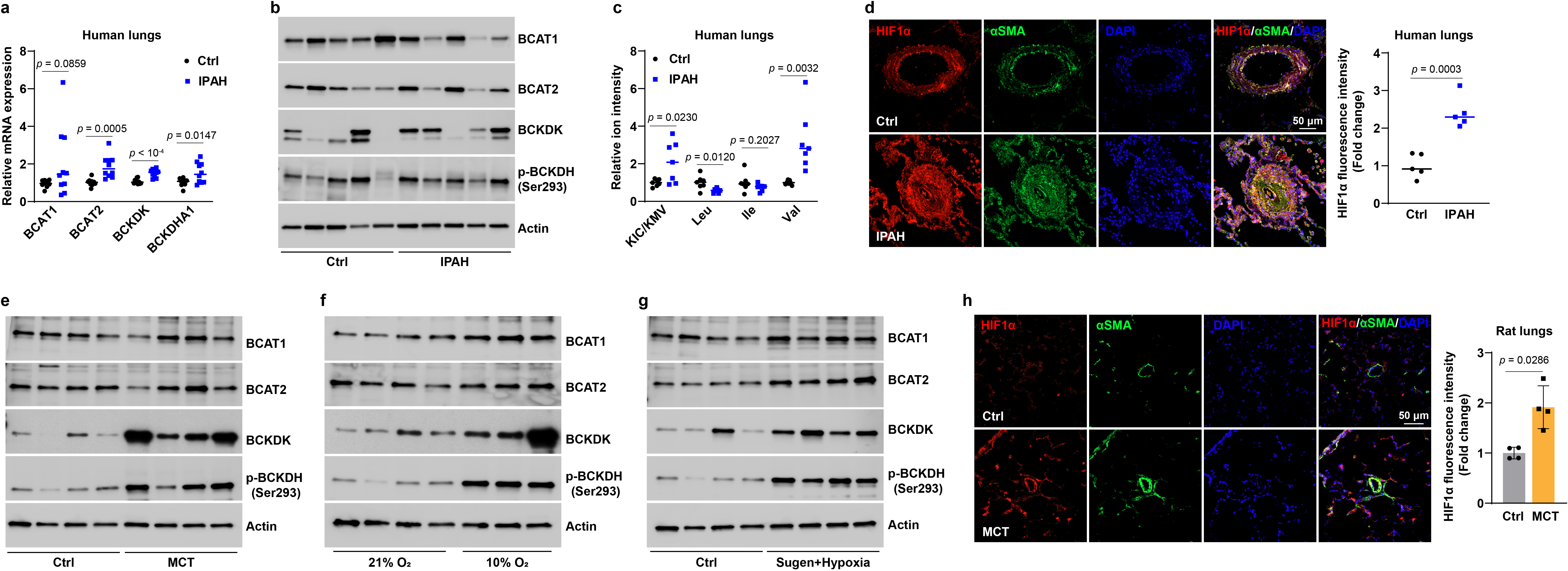
Dysregulated BCKA metabolism in PAH patients and animal models. **a,b,** mRNA (**a**) and protein (**b**) expression of key BCKA metabolic genes in human lung tissues of idiopathic PAH (IPAH) patients and transplantation-failed donors (Ctrl). Fold change was calculated relative to control donors. *n* = 10 individuals (**a**) and 8 individuals (**b**). **c,** The levels of BCKAs and BCAAs in the lungs of IPAH patients and control donors. Fold change was calculated relative to control donors. *n* = 7 individuals. **d,** Immunofluorescent images and quantitation of HIF1α protein in lung tissues of IPAH patients and control donors. Fold change was calculated relative to control donors. *n* = 5 individuals. **e-g,** Protein levels of four BCKA metabolic enzymes in lung tissues of control and PAH rats induced by MCT (**e**), hypoxia (10% O_2_, **f**), or Sugen5416 and hypoxia treatments (**g**). *n* = 4 (**e**, **g**) and 3-4 (**f**) rats per condition. **h,** Immunofluorescence images and quantitation of HIF1α protein in control and MCT-treated rat lung tissues. *n* = 4 rats. All data are presented as mean ± SD. Student’s t test (**a**, **c**, **d**) or Mann-Whitney U test (**h**) was conducted when compared to lungs from failed donors (**a**, **c**, **d**) or control rats (**h**).

We further characterized BCKA metabolism and HIF1α expression using three experimental PAH models. Intriguingly, elevated protein levels of BCKDK and p-BCKDH were observed in lung tissue of rats in which PAH is induced by monocrotaline (MCT), while BCAT1/2 protein levels were not affected (**Fig. 7e** and Extended Data Fig. 7b). Furthermore, we also noted increased levels of BCAT1, BCAT2, BCKDK, or/and p-BCKDH proteins in the lungs of two other distinct PAH rat models (hypoxia and Sugen5416 + hypoxia) (**Fig. 7f, g** and Extended Data Fig. 7c, d). In line with the upregulated HIF1α protein in VSMCs of IPAH lungs, a 2-fold increase in HIF1α protein was also observed in the VSMC layer of lungs from rats treated with MCT (**Fig. 7h**). These findings suggest that BCKA metabolism is dysfunctional in PAH patients and animals, and, as reported^34^, each of these three animal models partially recapitulates the gene expression patterns of the human disease.

We next explored the contribution of PASMCs to these genetic changes in lung tissue. Remarkably, PASMCs from IPAH patients (IPAH-PASMCs) expressed higher levels of BCAT2, BCKDK, p-BCKDH, and BCKDHA1, as well as lower levels of BCAT1, than did control PASMCs (**Fig. 8a, b**). These expression profiles were consistent with the results in lung tissue, indicating that PASMCs predominantly contributed to the changes of BCKA metabolic genes in PAH lungs.

**Fig 8.**
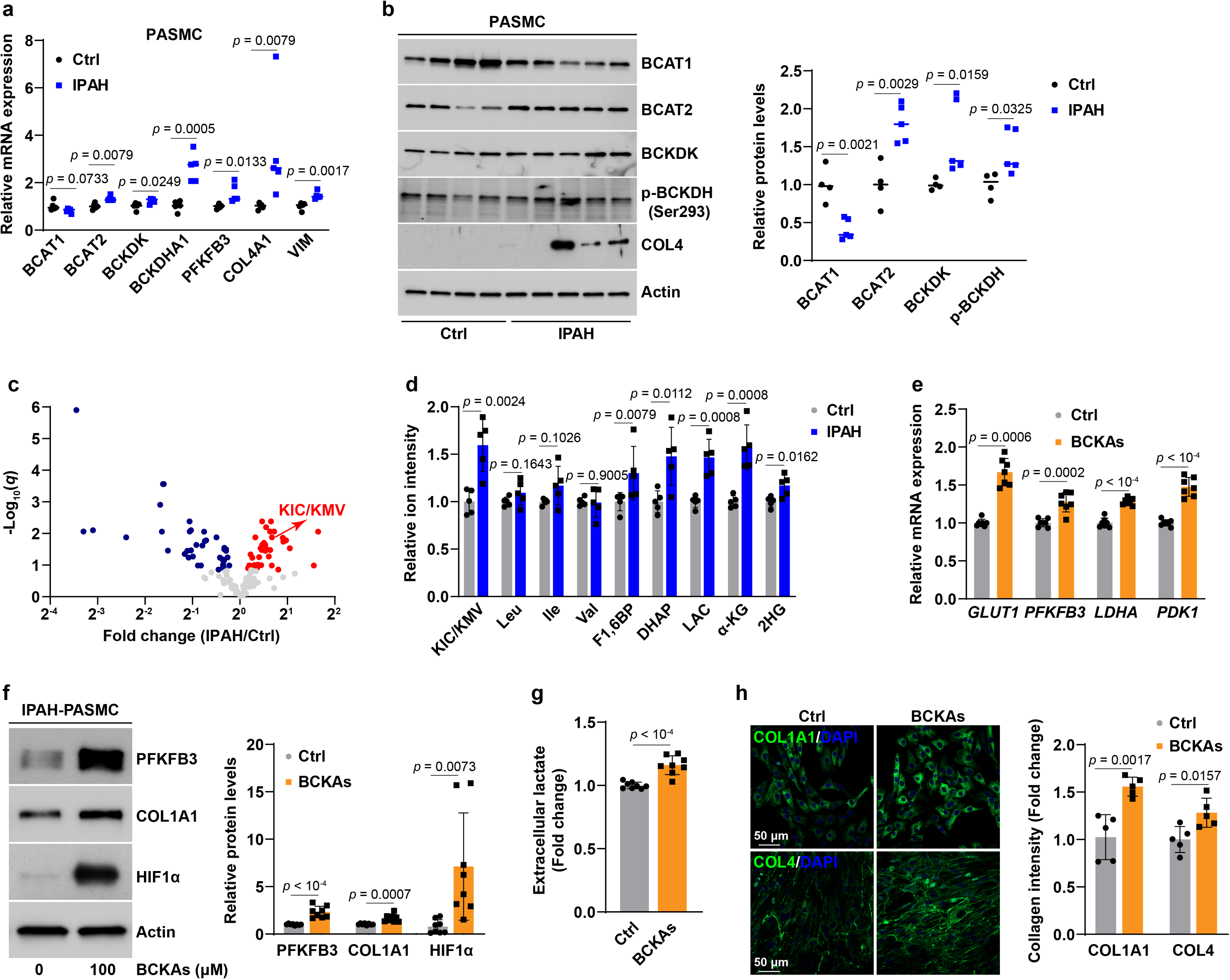
BCKA accumulation leads to enhancement of glycolytic activity and a synthetic phenotype in IPAH-PASMCs. **a,** mRNA expression of four BCKA metabolic enzymes, glycolytic gene *PFKFB3*, and synthetic markers in PASMCs from IPAH patients and commercially available control donors. Fold change was relative to control donors. *n* = 5 individuals. **b,** Protein levels of four BCKA metabolic enzymes and COL4 in PASMCs from IPAH and commercially available control donors. Fold change was relative to control donors. *n* = 4-5 individuals. **c,** Volcano plot from LC-MS metabolomic profiling showing the levels of 139 metabolites in normal and IPAH PASMCs. *n* = 5 individuals. **d,** LC-MS measurements of KIC/KMV, BCAAs, and glycolytic metabolites in normal and IPAH PASMCs. Fold change was calculated relative to normal PASMCs. *n* = 5 individuals. **e,f,** mRNA expression of HIF1α transcriptional target genes in glucose metabolism (**e**) and protein expression of HIF1α, PFKFB3, and COL1A1 (**f**) in BCKA-treated IPAH-PASMCs. Fold change was calculated relative to untreated IPAH-PASMCs. *n* = 7 (**e**) and 8 (**f**) from 4 individuals. **g,** Lactate secretion by IPAH-PASMCs treated with or without BCKAs. Fold change was calculated relative to untreated IPAH-PASMCs. *n* = 8 from 4 individuals. **h,** Confocal microscopy and quantitation results showing collagen synthesis and deposition in IPAH-PASMCs treated with vehicle control or BCKAs. Fold change was calculated relative to untreated IPAH-PASMCs. *n* = 5 from 4 individuals. All data are presented as mean ± SD. Student’s t test or Mann-Whitney U test was used when compared to PASMCs from normal control donors (**a**, **b**, **d**) or untreated PASMCs from IPAH patients (**e-h**).

Furthermore, metabolomic profiling revealed that KIC/KMV levels were 1.6-fold higher in IPAH-PASMCs than in normal controls, albeit their BCAA levels were comparable (**Fig. 8c, d**). Such increase in BCKA levels was close to the 2-fold elevation of KIC/KMV levels in IPAH lungs (**Fig. 7c**), implying that PASMCs are the major sources of BCKA accumulation. The accumulation of BCKAs correlated with induction of the key glycolytic enzyme *PFKFB3* and synthetic marker genes *VIM*, *COL4A1*, and COL4, and with elevations of glycolytic metabolites fructose 1,6-bisphosphate (F1,6BP), DHAP, and lactate levels, as well as α-KG and total 2HG levels in IPAH-PASMCs (**Fig. 8a-d**).

Such correlation was further substantiated by the effect of exogenous addition of BCKAs to cultured IPAH-PASMCs. In agreement with the results in normal PASMCs (**Figs. 3-5**), BCKAs activated HIF1α signaling under normoxia as evidenced by HIF1α protein stabilization, upregulation of its responsive genes GLUT1, PFKFB3, LDHA, and PDK1, and an increase in lactate secretion, supporting the conclusion that BCKAs enhanced glycolytic activity in IPAH-PASMCs (**Fig. 8e-g**). Furthermore, BCKAs also induced COL1A1 and COL4 gene expression as well as collagen synthesis, indicating promotion of synthetic phenotype (**Fig. 8f, h**). Silencing *BCAT1* and *BCKDHA1* significantly induced *COL1A1* and *COL4A1* mRNA expression and further heightened BCKA-induced upregulation of these two transcripts in IPAH-PASMCs; whereas the induction of *VIM* mRNA expression by BCKAs was attenuated by knockdown of *BCAT2* and *BCKDK*, albeit *COL1A1* and *COL4A1* mRNA expression did not change (Extended Data Fig. 8a-c). Taken together, these data demonstrate that dysregulated metabolism of BCKAs in PAH results in intracellular BCKA accumulation, and consequent promotion of enhanced glycolytic activity and synthetic phenotype in IPAH-PASMCs, both of which contribute to the pathobiology of this disease.

## Discussion

We here discover that intrinsic BCKAs, as newly identified PHD2 inhibitors, aerobically activate HIF1α signaling in normal VSMCs and provide the first mechanistic evidence linking BCKA metabolic dysregulation, aerobic HIF1α activity, and VSMC phenotypic dysfunction. Aerobic activation of HIF1α signaling has been well-documented in malignant cells^1,5–7,10,11,35^, while much less information is known about this mechanism in normal vascular cells. We show that HIF1α protein was detected in primary human VSMCs from pulmonary artery, aorta, and coronary artery, human placental pericytes, as well as Caco2 cancer cells under aerobic and unstimulated conditions, while it was not detectable in human micro/macrovascular endothelial cells and many other normal as well as cancer cells tested, supporting a cell type-specific and potentially paracrine phenomenon. In line with our findings, aerobic HIF1α protein stabilization was previously noted in human PASMCs but not in vascular endothelial cells^36–40^. However, these studies were uniformly interested in examining how external stimuli stabilized HIF1α, and none paid close attention to the observation of intrinsic HIF1α activation in aerobic cultures. To our knowledge, we are the first to elucidate the biological consequences and underlying mechanisms of paracrine stabilization of HIF1α protein by intrinsic factors under unstimulated and aerobic conditions. In addition to VSMCs, aerobic HIF1α activation has also been described in disturbed flow-stimulated human AoECs^41^, cell-permeable L2HG-treated HUVECs^27^, lipopolysaccharide (LPS)-challenged mouse macrophages^23,42^, KGDH or lipoic acid synthase depleted human dermal fibroblasts^10^, and skeletal muscle of ageing mice as well as *Sirtuin 1-*depleted mice^43^. Thus, these lines of evidence highlight the importance of understanding aerobic HIF1α activation in normal primary cells and in the setting of organism homeostasis.

Thus far, there are two well-studied mechanisms responsible for aerobic HIF1α stabilization: induction of ROS and competition with α-KG. Compelling evidence from primary VSMC studies supported that cellular ROS, particularly hydrogen peroxide (H_2_O_2_), activated HIF1α signaling through transcriptional and posttranslational mechanisms^37,44–48^. Theoretically, the catalytic Fe^2+^ center of PHDs can also be inactivated by H_2_O_2_ *via* Fenton chemistry and by superoxide *via* direct oxidization leading to stabilization of HIF1α. We did not observe any significant changes in cellular ROS levels of PASMCs cultured in CM, ruling out the possibility of ROS being the principal mechanism of aerobic HIF1α activation in our experimental conditions. The second known mechanism is the competitive displacement of α-KG bound to PHDs. This action was mostly carried out by the TCA cycle intermediary metabolites that have structural similarities with α-KG, such as fumarate, succinate, and malate^4,5,7,8^. Interestingly, we also excluded these known metabolites being the key mediators of paracrine activation of HIF1α in vascular cells in normoxia. Instead, we identified BCKAs as the mediators of paracrine activation of HIF1α signaling in vascular cells under aerobic conditions, a phenomenon that has not been previously described, despite the fact that other mechanisms, such as decreased iron incorporation into PHD2 protein^49^, deubiquitination, and phosphorylation of HIF1α^50,51^, may be also involved in aerobic stabilization of HIF1α protein and warrant future investigation.

More importantly, we provided compelling and novel mechanistic evidence for the mechanism by which BCKAs stabilize HIF1α under well-oxygenated conditions. We discovered that BCKAs inhibit PHD2 activity through direct and indirect mechanisms. First, these metabolites directly suppress PHD2 hydroxylation activity likely by chelating its catalytic Fe^2+^ active center with an IC_50_ value of 252 μM (higher than that of currently developed specific, potent inhibitor roxadustat IC_50_ = 17 μM), indicating BCKAs are relatively weak PHD2 inhibitors, albeit in a physiologically relevant range. Nevertheless, our discoveries suggest that BCKAs may serve as genetic and epigenetic modifiers by inhibiting the activity of other dioxygenases, such as collagen prolyl 4-hydroxylases, JmjC-domain-containing histone lysine demethylases, and the ten-eleven translocation DNA demethylases. Furthermore, as plasma BCKA levels in maple syrup urine disease patients reached 1-4 mM^20^, it is of great interest to investigate the effects of BCKA accumulation on the activities of HIF1α, PHDs, and other dioxygenases in this rare disease patient population.

The second mechanism responsible for BCKA-induced PHD2 inhibition is BCAT1-catalyzed production of α-KG coupled with LDHA-mediated generation of L2HG. We provided consistent and compelling evidence for this unique mechanism by modulating L2HG metabolic enzymes and exogenous supplementation of L2HG. Discovery of this additional mechanism strongly strengthened our hypothesis that BCKAs stabilize HIF1α in normoxia. Previous studies showed that KMV when used at 2-20 mM increased intracellular α-KG levels by inhibition of KGDH activity^22,27,28,52,53^. Here, we found that an increase of α-KG production could be specifically achieved by BCAT1-mediated reamination of BCKAs when used at 50-200 μM concentrations, highlighting a novel mechanism of physiological relevance. L2HG has been well-described as a potent PHD2 inhibitor in cancer and normal cells^10,23–25,27,35^. The enzymes responsible for L2HG accumulation are of great importance here. Williams *et al*.^23^ found that suppression of L2HGDH expression was responsible for LPS-induced L2HG accumulation in macrophages. Recent work revealed that KMV-mediated inhibition of KGDH activity resulted in increased L2HG production in quiescent HUVECs with *forkhead box O1* overexpression^27^. Both studies did not examine the roles of LDHA and MDH1/2, two well-known enzymes that generate L2HG under hypoxia and acidic pH in normal and malignant cells^21,22,35^. In aerobic PASMCs, we discovered that the key enzyme for L2HG accumulation is LDHA, consistent with the observations showing LDHA but not MDH1/2 produced L2HG in hypoxic CD8^+^ T lymphocytes^54^. LDHA-mediated production of lactate is expected to lower cellular pH. An acidic pH (*e.g.,* pH = 6.0∼6.8) was able to increase total 2HG or L2HG levels by enhancement of LDH activity (with lesser effects on MDH activity), inhibition of L2HGDH activity, and promotion of the binding of protonated α-KG with LDHA in aerobic cultures of glioblastoma and HEK293 cells^28,35^. It remains unclear as to whether and to what extent this mechanism contributes to L2HG production in our study. Furthermore, as known under hypoxic HIF1α activation^54^, we also found that LDHA basal expression and BCKA-induced LDHA upregulation were HIF1α dependent under normoxia. Thus, our findings reveal a positive feedforward mechanistic loop, where BCKAs activate HIF1α to stimulate LDHA expression, which then enhances L2HG production to inhibit PHD2 activity, as a result further enhancing HIF1α activity under aerobic conditions.

It is noteworthy that BCKA-induced aerobic HIFα activation has unique cell type and isoform specificity. One common feature shared by these responsive cell types was that HIF1α protein was easily detected under basal and aerobic conditions, while it was absent in the nonresponsive cells, further confirming that BCKAs are the paracrine stabilizers of HIF1α. The underlying mechanisms for such discrepancy remain to be determined; but may include changes in expression patterns of BCKA metabolic enzymes; differences in intracellular α-KG and L2HG levels; changes in the expression levels of L2HG metabolic enzymes LDHA; MDH1/2, and L2HGDH; and changes in the expression levels of PHD proteins. Furthermore, our findings that inhibition of PHD2 activity by BCKAs led to selective induction of HIF1α but not HIF2α protein are also of great interest. This result could, in part, be a consequence of the greater abundance of *HIF1α* mRNA as compared to *HIF2α* in VSMCs, and or a consequence of PHD2 preferentially hydroxylates by PHD2 of HIF1α protein to regulate its stability^55–57^, in part, owing to HIF-associated factor-selective binding to and (further) stabilization of HIF1α^16^ and of PHD3 induction by HIF1α destabilizing HIF2α^17^.

As the functional consequences of BCKA-induced aerobic HIF1α activation, we focused on glucose metabolism and modulation of a phenotypic switch in PASMCs. We first observed that glycolytic activity was enhanced by BCKAs in a HIF1α-dependent manner, which validated the canonical response of HIF1α activation in mammalian cells under both adequate and low oxygen conditions^4,23,41^. Surprisingly, we also found that BCKAs promote phenotype switching toward a synthetic subtype in PASMCs, most strikingly evidenced by increased collagen gene expression and biosynthesis. The mechanisms governing this phenotypic change remain unclear. The phenotypic plasticity of VSMCs is controlled by a complex of genetic, epigenetic, mechanical, and metabolic mechanisms^29,31^. A recent study showed that supplementation of lactate in growth medium promoted a synthetic phenotype in human induced pluripotent stem cell derived VSMCs^58^. Under hypoxia, HIF1α-mediated microRNA-9 upregulation correlated with a synthetic phenotype in rat PASMCs^32^. By silencing *HIF1α*, we here noted that BCKA-induced increases in lactate levels were abolished with concomitant enhancement of collagen synthesis and increased contraction of PASMCs in collagen gels, indicating that BCKAs *per se* promote collagen synthesis independent of lactate production and that BCKA-induced HIF1α activation, in turn, functions as a negative feedback mechanism in this specific cell type. Indeed, Barnes *et al*^39^ reported that *HIF1α* silencing also increased collagen gel contraction in IPAH-PASMCs by enhancing myosin light chain phosphorylation. By contrast, loss of *Hif1α* abrogated angiotensin II-induced upregulation of *Col1a1* and *Col3a1* mRNA in mouse AoSMCs *ex vivo*^59^. These lines of evidence highlight the distinct and complex roles of HIF1α in regulating VSMC functions in different vascular beds.

The elevated levels of circulating BCKAs and more notably BCAAs have been positively linked to many cardiovascular metabolic diseases^19,26,60–63^. Proposed mechanisms include mitochondrial metabolic and oxidative stress, activation of mTOR and MEK-ERK signaling pathways, suppression of insulin signaling pathways, platelet activation, and inflammation^19,26,60–63^. We here showed that the expression of four key BCKA metabolic enzymes, BCAT2, BCKDK, BCKDHA1, and p-BCKDH, were mostly upregulated to different extents in PAH lungs and PASMCs, leading to BCKA accumulation and, as a result HIF1α activation, enhanced glycolysis and increased collagen synthesis, all of which are observed in PAH pathobiology^15,30,33^. We believe that this is the first piece of evidence showing a correlation between dysregulation of BCKA metabolism specifically in PASMCs and PAH phenotypes. However, how dysregulation of the metabolism of BCKAs *per se*, BCKA-induced HIF1α activation, and BCKA-induced collagen biosynthesis in PASMCs as well as other pulmonary vascular cells contribute to the development of PAH warrants future investigation.

In summary, we conclude that BCKAs inhibit PHD2 activity to activate HIF1α signaling, enhance glycolytic activity, and promote a synthetic phenotypic switch in PASMCs, which together contribute to vascular dysfunction, including the development of PAH.

## Methods

### Chemicals and reagents

Sodium salt of 3-methyl-2-oxobutanoic acid (KIV; cat # AC189720050) was purchased from Thermo Fisher Scientific (Waltham, MA). Sodium salt and acid forms of 4-methyl-2-oxovalerate (KIC; cat # K0629-1G and 68255-1G), 3-methyl-2-oxovaleric acid sodium salt (KMV; cat # K7125-5G), sodium butyrate (cat # B5887-1G), 3-hydroxyisobutyrate (cat # 36105-250MG), β-hydroxybutyrate (cat # 21822), sodium lactate (cat # 1614308), sodium pyruvate (cat # P4562), MG132 (cat # M7449-200UL), proteinase K (cat # P2308), monocrotaline (MCT; cat # C2401-1G), and other unspecified chemicals were purchased from Sigma Chemical Co. (St. Louis, MO). Cell permeable trifluoromethylbenzyl esters of L-2-hydroxylglutarate (L2HG) and its enantiomer D2HG were synthesized by WuXi App Tec (Shanghai, China). Recombinant human transforming growth factor β (TGFβ; cat # 100-21) and platelet-derived growth factor (PDGF-BB; cat # 220-BB-010) were obtained from PeproTech (Cranbury, NJ) and R&D Systems (Minneapolis, MN), respectively.

Sugen5416 (cat # A3847) was from APExBIO (Houston, TX). DMEM without amino acids was purchased from US Biological (Cat # D9800-13; Salem, MA).

### Cell culture and treatments

All primary cells were obtained commercially. Primary human pulmonary artery smooth muscle cells (PASMCs; cat # CC-2581) and endothelial cells (PAECs; cat # CC-2530), human aortic smooth muscle cells (AoSMCs; cat # CC-2571) and endothelial cells (AoECs; cat # CC-2535), human coronary artery smooth muscle cells (CASMCs; cat # CC-2583) and endothelial cells (CAECs; cat # CC-2585), human umbilical vein endothelial cells (HUVECs; cat # C2517A), human lung microvascular endothelial cells (MVECs; cat # CC-2527), human lung fibroblasts (LFs; cat # CC-2512), and human normal dermal fibroblasts (NDFs; cat # CC-2511) were purchased from Lonza (Walkersville, MD). Human THP1 monocytes (cat # TIB-202), HeLa cells (cat # CRM-CCL2), and rat H9c2 cardiomyocytes (cat # CRL-1446) were obtained from ATCC (Rockville, MD). Human placental pericytes were purchased from PromoCell (cat # C-12980; Heidelberg, Germany). Human cancer lines skin squamous cell carcinoma IC8 (IC8SCC) and breast epithelial carcinoma MDA-MB157 cells (Jon C. Aster), lung epithelial carcinoma H1299 and H661 cells (Li Chai), colorectal adenocarcinoma Caco2 cells (Richard S. Blumberg), and hepatocellular carcinoma HepG2 cells (David Cohen) were kindly provided by our colleagues in Departments of Medicine and Pathology at Brigham and Women’s Hospital. For SMCs, passages 3 to 5 were used in all experiments; for endothelial cells and other normal cell types, passages 3 to 8 were used; and for cancer lines, up to 8 passages were used in experiments.

PASMCs, AoSMCs, CASMCs, and IPAH-PASMCs were cultured in SmBM-2 medium (cat # CC-3181) with SingleQuots supplements (cat # CC-4149); macrovascular (PAECs, AoECs, and HUVECs) and microvascular (CAECs and MVECs) endothelial cells were grown in EBM-2 medium (cat # CC-3156) supplemented with EGM-2 (cat # CC-4176) or EGM-2MV (cat # CC-4147) BulletKit growth factors, respectively; pericytes were also grown in EBM-2 medium with the addition of EGM-2 BulletKit growth factors without ascorbate; and LFs and NDFs were grown in FGM-2 growth medium (cat # CC-3131) with supplementation of growth factors (cat # CC-4126). All media for primary cells were purchased from Lonza. High glucose DMEM (cat # 11965092; Thermo Fisher Scientific, Waltham, MA) supplemented with 10% fetal bovine serum was used to grow THP1, H9c2, MB157, HeLa, HepG2, and IC8SCC cells; and complete RPMI-1640 medium (cat # 11875119; Thermo Fisher Scientific) was used to culture THP1, H1299, and H661 cells.

All cells were maintained at 37°C in an incubator with 95% humidity and 5% CO_2_. Medium was refreshed every other day for cell growth and immediately prior to conditioned medium (CM) or chemicals treatment. An equal volume of ultrapure DNase/RNase-free distilled water or sterile DMSO was used as vehicle control. For hypoxic experiments, cells were exposed to 0.2% O_2_ for 24 hours using a modular hypoxia chamber.

### Metabolomic profiling and analysis

#### Sample preparation

PASMCs were quickly washed with two volumes of ice-cold PBS, lysed for 15 min in pre-chilled 80% liquid chromatography-mass spectrometry (LC-MS)-grade methanol, and harvested by scraping on dry ice followed by centrifugation at maximal speed for 30 min at 4°C. For medium samples, 50 μL of medium were incubated in 200 μL of pre-cooled 100% LC-MS-grade methanol for 15 min on dry ice and then centrifuged as done for cell samples. The supernatants of cell and medium samples were collected and evaporated to dryness on an Integrated SpeedVac System (Thermo Fisher Scientific; Waltham, MA) at 42°C. The resulting pellets were resuspended in 50 μL of LC-MS-grade water. Twenty microliters of sample were loaded in an autosampler vial for analysis.

#### Mass spectrometric acquisition

LC-MS analysis was performed on a Vanquish ultra-high performance liquid chromatography system coupled to a Q Exactive mass spectrometer (Thermo Fisher Scientific) that was equipped with an Ion Max source and HESI II probe using the protocol described previously^22,64^. Briefly, metabolites were separated using a ZIC-pHILIC stationary phase (150 mm × 2.1 mm × 3.5 mm; Merck) with a guard column. Mobile phase A contained 20 mM ammonium carbonate and 0.1% ammonium hydroxide. Mobile phase B was acetonitrile. The mobile phase flow rate was 100 μL/min and gradient (%B) was 0 min, 80%; 20 min, 20%; 20.5 min, 80%; 28 min, 80%; and 42 min, stop. The column effluent was introduced into the mass spectrometer with the following ionization source settings: sheath gas 40, auxiliary gas 15, sweep gas 1, spray voltage +3.0 or -3.1 kV, capillary temperature 275°C, S-lens RF level 40, and probe temperature 350°C. The mass spectrometer was operated in polarity-switching full scan mode from 70-1000 m/*z*. Resolution was set to 70,000, and the AGC target was 1×10^6^ ions. Data were acquired and analyzed using TraceFinder 4.1 (Thermo) with peak identifications based on an in-house library of authentic metabolite standards previously analyzed utilizing this method. Missing values were imputed using random forest. Sample peak areas were normalized using probabilistic quotient normalization^65^.

#### Mass spectrometry data analysis

The normalized data were analyzed using MetaboAnalyst 5.0^66,67^. The statistical analysis module was first applied to test the statistical significance of metabolite differences between untreated and treated groups. Based upon the initial statistical analysis, a list of metabolites with significant changes was further analyzed using metabolite-set enrichment analysis (MESA) module to identify biological pathways that were enriched by treatments.

### [^13^C_6_]-KIC isotope tracing

PASMCs were seeded into a 6-well plate and labeled in complete fresh SmBM-2 medium supplemented with 1 mM uniformly [^13^C]-labeled KIC (^13^C_6_-KIC) isotope (cat # CLM-4785-0.1, Cambridge Isotope Laboratories; Tewksbury, MA) for 8 hours. Metabolites were extracted and analyzed by LC-MS as described above with the following modifications. To increase sensitivity for specific metabolites of interest and their isotopes, the mass spectrometer was operated in selected ion monitoring mode using an m/z window of 9 centered on the range of isotopes for a given molecule. Raw peak areas were corrected for quadrupole bias^68^, and the resulting mass isotopomer distributions (MIDs) were corrected for natural isotope abundance using a custom R package encoding the method of Fernandez *et al*^69^.

### High-performance liquid chromatography (HPLC) measurements of BCKAs

BCKAs levels in medium were determined by HPLC-fluorescence detection as previously published^70,71^ with minor modifications. Medium samples were centrifuged at 1,200 rpm for 5 min at 4°C to remove cell debris. One hundred eighty microliters of medium were mixed with 20 μL of internal standard 2-oxovaleric acid (1 mM; cat # 75950-50 mL) and deproteinized by acetonitrile followed by centrifugation at 12,000 rpm for 10 min at 4°C. The supernatants were evaporated to approximately the starting volume 200 μL on an Integrated SpeedVac System. For derivatization, forty microliters of medium were mixed with an equal volume of derivatization compound O-phenylenediamide (OPD; cat # P23938-5G; 25 mM in 3 N HCl) and then heated at 80°C for 30 min. Twenty microliters of the resulting supernatants were loaded to an autosampler vial for fluorescence detection (excitation at 350 nm, emission at 410 nm) using a C18 column on a 1260 Infinity II LC System (Agilent; Lexington, MA). The mobile phase solutions A and B were 0.1% formic acid in LC-MS-grade water and acetonitrile, respectively. The gradient conditions were as follows: t = 0 min, 95% A and 5% B; t = 11 min, 40% A and 60% B; t = 11.10 min, 100% B; and t = 14.10 min, 100% B; the flow rate was set at 0.42 mL/min. BCKA standards were processed exactly as medium samples and a standard curve was generated to calculate the concentrations of BCKAs.

### Extracellular metabolic flux analysis

Extracellular acidification rate (ECAR) and oxygen consumption rate (OCR) were analyzed using the Seahorse extracellular flux (XF) analyzer (Agilent; Lexington, MA) as we previously described^64,72^. Ten thousand cells were seeded on a Seahorse microplate in 250 μL growth medium. After overnight attachment, cells were cultured in CM or treated with BCKAs for 8 hours. For the glycolysis stress test, XF base medium (cat # 103335-100; Agilent) was supplemented with 2 mM glutamine and pH was adjusted to 7.4. ECAR and OCR were recorded at baseline and followed by sequential addition of glucose (final concentration 10 mM), of oligomycin (final concentration 1 μM), and of 2-deoxyglucose (final concentration 50 mM). Data were acquired and analyzed using Wave software (Agilent). Cell counts in each well were determined using a Beckman Coulter counter and used to normalize flux rates.

### Quantitation of extracellular lactate

Secreted lactate levels were measured by a fluorescence-based assay kit (cat # 700510, Cayman Chemical; Ann Arbor, MI). Cell culture medium was collected and centrifuged (1,200 rpm, 5 min at 4°C) to remove cell debris. The resulting medium was used for the assay according to the manufacturer’s protocol. Fluorescence was measured using a SpectraMax i3x microplate reader (Molecular Devices; San Jose, CA). Lactate concentration was calculated based upon a standard curve of L-lactate. The amount of lactate was calculated by multiplying its concentration with the initial sample volume and dilution factor, then normalized to total protein content for each sample. Fold change was calculated relative to control cells under normoxia.

### Determination of PHD2 enzymatic activity

PHD2 activity was measured by the determination of α-KG consumption using a colorimetric assay as recently published^73^. For IC_50_ experiments, each reaction was conducted in individual wells of a 384-well plate in a final volume of 30 µL. KIC, KIV, KMV, BCKAs, and Roxadustat (a known PHD2 inhibitor; cat # HY-13426, MedChem Express; Monmouth Junction, NJ) were titrated in their respective concentrations into 384-well plates using a D300e Dispenser. Fifteen microliters of MBP-HA-PHD2 enzyme (1.4 mg/mL) was dispensed into each well and incubated with the compounds for 10 min at room temperature, followed by addition of 15 µL of cofactor mix to start the reactions [final concentrations: 50 mM MES (pH = 6.0), 0.6 mg/mL catalase, 1 mM dithiothreitol (cat # DTT10, GoldBio; St Louis, MO), 500 µM ascorbic acid, 50 µM FeSO_4_, 250 µM α-KG, and 100 µM of HIF1α peptide (GenScript; Piscataway, NJ)]. The reactions were allowed to proceed at room temperature to prevent uneven plate heating that could affect reaction rates. At time points 0, 10, 20, 30, 45, and 90 min, respective wells were quenched with 30 µL of 10% trichloroacetic acid. Plates were then centrifuged at 1,500 rpm for 15 min, and 30 µL of clarified supernatants transferred to new 384-well plates using a Bravo automated liquid handling platform (Agilent) and incubated with an equal volume of 50 mM 2,4-dinitrophenylhydrazine (2,4-DNPH) solution for 30 min at room temperature to derivatize α-KG, followed by 15 µL of 6 M sodium hydroxide for 10 min at room temperature before reading plates at 420 nm on an EnVision plate reader (PerkinElmer; Waltham, MA). Initial velocities for each inhibitory BCKAs concentration were derived by regression fitting of timepoints within distinct linear ranges of each time course. IC_50_ values were derived from curves were fitted to a log(inhibitor) versus response function in GraphPad Prism 9.0.

### Simulation of molecule docking

Molecular Operating Environment (MOE; Montreal, QC, Canada), an integrated computer-aided molecular design platform, was used to predict the potential binding sites and estimate the docking energy of KIC, KIV, and KMV to the PHD2 enzyme (tertiary structure accession ID: 2G19). The protein-ligand interaction fingerprint (PLIF) module was used to map various potential binding configurations of each BCKA with the PHD2 active site. The two known binding ligands of PHD2, α-KG and roxadustat, were included for reference.

### Determination of L2HG levels

Cellular L2HG levels were specifically determined using an enzymatic assay we developed recently^74^. The assay is based on the oxidation of L2HG to α-KG coupled to the reduction of FAD to FADH_2_ in the presence of L2HG dehydrogenase (L2HGDH); the intermediate is further converted to pyruvate and hydrogen peroxide by enzyme mixture alanine transaminase, pyruvate oxidase, and horseradish peroxidase, which reduces 10-acetyl-3,7-dihydroxypheno-xazine to a colored product resorufin with strong absorbance at 570 nm. This absorbance is closely proportional to the amount of L2HG present in the samples.

Cell extracts were harvested by scraping in pre-cooled 80% LC-MS-grade methanol and centrifuged at maximal speed for 10 min at 4°C. The obtained supernatants were transferred to a new tube and evaporated to dryness using a SpeedVac. In order to minimize the influence of the endogenous L2HGDH and other dehydrogenases, the obtained cell powder was reconstituted in 100 μL of 0.6 N PCA and incubated on ice for 1 hour followed by addition of neutralizing buffer for pH adjustment to 7.0. The resulting solution was centrifuged at 14,000 g for 10 min at 4°C to remove the denatured endogenous dehydrogenases. The samples were then reconstituted in 200 μL of the assay buffer containing 50 mM HEPES, 5.0 mM L-alanine, 1.2 mM MgSO_4_, 1 mM TPP, 0.2 mM FAD, 4.7 mM KCl, 1.2 mM KH_2_PO_4_, 118 mM NaCl, and 2.5 mM CaCl_2_ (pH = 7.4). The reactions took place at 37°C for 2 hours in duplicates in the presence or absence of L2HGDH. The assay was initiated by addition of L2HGDH positive enzyme mixture (containing 0.2 U alanine transaminase, 0.2 U pyruvate oxidase, 0.2 U horseradish peroxidase, and 40 nmol 10-acetyl-3,7 dihydroxyphenoxazine, and 10 μg L2HGDH) or L2HGDH negative enzyme mixture (containing all enzymes in positive mixture except L2HGDH, but with equivalent bovine serum albumin instead). A blank in the absence of substrate and enzymes was included. The absorbance readings in the blank and in the L2HGDH negative mixture were subtracted from the readings in the L2HGDH positive mixture. The L2HG content in samples were calculated from a L2HG standard curve and normalized to total protein content of samples determined by the Lowry method.

### Medium conditioning and treatments

CM was harvested from PASMCs cultures after 24 hours of incubation under aerobic conditions, centrifuged to remove cell debris, sterilized using 0.22 μm filters, and stored at -80°C until use. For heat inactivation of secreted proteins in medium, complete VSMCs growth medium (GM) and CM were heated up in 95°C water bath for 30 min. GM and CM were also incubated with proteinase K (50 µg/mL) in 37°C water bath for 2 hours followed by incubation at 95°C for 20 min to inactivate any residual activity of proteinase K. For medium fractionation, a centrifugal filter with 10 kDa cutoff size (cat # UFC901008, EMD Millipore) was used to fractionate GM and CM by centrifugation at 5,000 g for 20 min at 4°C. The fraction larger than 10 kDa (∼500 μL) was then resuspended with the original volume of unfractionated medium. The resulting media from all three approaches were filtered through 0.22 µm syringe filter to for sterilization before reapplication.

### IPAH patient specimen

Human lung frozen tissues, slides, and RNA samples of IPAH patients and control donors as well as PASMCs of IPAH patients were obtained from the Pulmonary Hypertension Breakthrough Initiative (PHBI) under an approved protocol by the IRB committee at Brigham and Women’s Hospital (IRB protocol # 2021P001557). The clinical and demographical information of human subjects were available at Supplementary Table 1.

### PAH patient cohort

Peripheral venous blood samples were obtained from a total of 32 PAH patients at Brigham and Women’s Hospital. Patients underwent a clinically indicated right heart catheterization to measure hemodynamic parameters. Plasma derived from the whole blood samples were processed, and metabolites were analyzed using the Metabolon Global Metabolic HD4 platform. The data were log_2_-transformed and missing data were imputed and then normalized using a centered log-ratio method. The study was approved by the Massachusetts General Brigham Institutional Review Board (2016P001269) and all participants provided written informed consent for using their clinical data and biological specimen for research. The Spearman correlation coefficient was calculated to evaluate the associations between total BCAAs/total BCKAs ratio and mPAP or PVR.

### Animal models of PAH

PAH animal models were established using our previously well-described protocols^75,76^. We chose to use only male animals as we and others routinely do so to minimize the attenuating influences of sex on the pathobiology of the disease^77,78^. Animals were randomly allocated to different treatment groups. For MCT-induced PAH, adult Sprague-Dawley (SD) rats (12-14 weeks of age; Charles River Laboratory) were intraperitoneally injected a single dose of MCT (60 mg/kg BW) and sacrificed 3 weeks later. For hypoxia-induced PAH, adult SD rats were housed in 10% normobaric oxygen environment for 21 days. For sugen5416 and hypoxia-induced PAH, adult SD rats were subcutaneously injected once with sugen5416 (20 mg/kg BW; a VEGFR-2 kinase inhibitor) and then placed in 10% normobaric oxygen environment for 3 weeks. Control animals received vehicle solvent injection and housed in 21% normobaric oxygen condition for 3 weeks. The lung tissues were snap-frozen in liquid nitrogen for gene expression analyses. All animals were housed at Brigham and Women’s Hospital Animal Facility with a 12:12-hour light:dark cycle, a 50-60% relative humidity, and *ad libitum* access to water and food. All animal procedures were approved by the IACUC at Brigham and Women’s Hospital.

### Three-dimensional collagen matrix contraction assay

The contractility of PASMCs was evaluated using a collagen matrix contraction assay (cat # CBA-201, Cell Biolabs; San Diego, CA). Briefly, PASMCs were mixed with ice-cold type I collagen gel working solution at a ratio of 2:8 (v/v), which were then dispensed to a 24-well plate and incubated at 37°C for 1 hour to allow collagen polymerization. After 8 hours of BCKA treatment, the collagen matrix gel was gently released from the sides of each well with a sterile 200 μL pipette tip. The plate images were captured at time 0 and 8 hours post gel detachment using a ChemiDoc Touch Imaging system (Bio-Rad) and quantitated using Image J v1.54 software (NIH; Baltimore, MD). The contracted area of collagen gel was calculated as the area differences between time 0- and 8-hour and then normalized to controls.

### Gene silencing by siRNAs

ON-TARGETplus siRNAs targeting human *HIF1α* (cat # L-004018-00-0005), *BCAT1* (cat # L-012084-00-0005), *BCAT2* (cat # L-020191-00-0005), *LDHA* (cat # L-008201-00-0005), *L2HGDH* (cat # L-008130-01-0005), *BCKDHA1* (cat # L-012563-00-0005), *BCKDK* (cat # L-004932-00-0005), or non-targeting control (cat # D-001810-10-05) were purchased from Dharmacon (Lafayette, CO). PASMCs were transfected with 5 pmol of each siRNA using Lipofectamine RNAiMAX transfection reagent (cat # 13778-150, Life Technologies; Carlsbad, CA). After overnight transfection, cell medium was refreshed for 24 hours, and cells were then cultured in CM or treated with BCKAs as indicated. The efficacy of gene silencing was confirmed by detection of mRNA and/or protein expression.

### Quantitative real time PCR

Total RNA was isolated using RNeasy Mini kit (cat # 74106, Qiagen; Germantown, MD) according to the manufacture’s protocol. The quality and concentration of purified RNA samples were evaluated using a Nanodrop One spectrophotometer (Thermo Fisher Scientific). One microgram of total RNA was reverse transcribed into cDNA in a 25 µL reaction system using a High Capacity cDNA Reverse Transcription kit (cat # 4368813, Applied Biosystems; Foster City, CA). Diluted cDNA was then used to amplify target genes using TaqMan primers and a fluorogenic detection probe on a 7900HT Fast Real-time PCR machine (Applied Biosystems) following initial steps of 50°C 2 min and 95°C 10 min, and then 40 cycles of 95°C 15 sec for denaturation and 60°C 1 min for annealing and extension. β-actin expression in each sample was used as an internal control. Gene expression was calculated using the 2^-ΔΔCt^ method.

### Immunoblotting

Cells were promptly rinsed with ice-cold PBS before harvesting and then lysed by sonication in RIPA buffer supplemented with proteinase and phosphatase inhibitors (cat # sc-24948, Santa Cruz Biotech; Santa Cruz, CA). Ten micrograms of total proteins were separated on 4-15% pre-cast SDS-PAGE gels and then transferred to PVDF membranes using a semi-dry Turbo transfer system (Bio-Rad; Hercules, CA). Protein blots were blocked in 5% blotting-grade blocker milk for 1 hour at room temperature, and then incubated with specific primary antibodies (1:1,000 dilution in 2% (w/v) BSA in TBS with 0.1% Tween-20) overnight at 4°C on a rotating platform. HRP-linked secondary mouse or rabbit antibody (Cell Signaling Technology; Danvers, MA; 1:5,000 dilution) and ECL detection reagents (cat # K-12045-D20, Advansta; Menlo park, CA) were employed to visualized protein blots. Images were captured using a ChemiDoc Touch Imaging system and quantitated by Image Lab software 5.2 (Bio-Rad).

Human BCAT1 (cat # 611271) and HIF1α (cat # 610958) antibodies were from BD Biosciences; human BCAT2 (cat # 9432), LDHA (cat # 2012), MDH2 (cat # 11908), PHD2 (cat # 3293), pVHL (cat # 68547), hydroxylated HIF1α-Pro564 (cat # 3434), VIM (cat # 5741), human and rat phospho-BCKDH-E1α (Ser293) (cat # 40368) antibodies were purchased from Cell Signaling Technologies; human COL1A1 (cat # NBP1-30054), COL4 (cat # NB120-6586), and HIF2α (cat # NB100-122) antibodies were obtained from Novus Biologicals; rat Bcat1 (cat # 13640-1-AP), rat Bcat2 (cat # 16417-1-AP), and human PFKFB3 (cat # 13763-1-AP) antibodies were purchased from Proteintech; human and rat BCKDK (cat # sc-374424) antibody was from Santa Cruz Biotech; human MDH1 (cat # ab175455), PDK1 (cat # ADI-KAP-PK112-D), and BCKDHA1 (cat # A303-790A) antibodies were from Abcam, Enzo Life Sciences, and Bethyl Laboratories, respectively. β-actin (cat # sc-47778, Santa Cruz Biotech) antibody was included as loading controls. Cellular protein concentration was determined using the DC protein assay (Bio-Rad).

### Immunofluorescence staining

Formalin-fixed and paraffin-embedded human lung tissue slides (5 µm thickness) were deparaffinized, rehydrated, and then boiled for 15 min in citric acid buffer (5 mM, pH = 6.0) for antigen retrieval. For immunocytochemistry assay, cells were seeded on 4-well Nunc Lab-Tek chamber slide and fixed with 4% paraformaldehyde for 10 min. Tissue and cell slides were then blocked with 1% BSA and 10% normal goat serum in PBS at room temperature for 1 hour followed by overnight incubation at 4°C with primary antibodies (1:50 dilution) against human HIF1α (cat # 610958, BD Biosciences), rat HIF1α (cat # NB100-105, Novus Biologicals), human smooth muscle actin alpha (αSMA; cat # ab124964, Abcam), rat αSMA (cat # 5694, Abcam), human COL1A1 (cat # NBP1-30054, Novus Biologicals), or human COL4 (cat # NB120-6586, Novus Biologicals), and then Alexa Fluor^®^ 568 goat anti-mouse (cat # ab175473, Abcam), 568 goat anti-rabbit (cat # 175471, Abcam), 488 goat anti-mouse (cat # 1500113, Abcam), or 488 goat anti-rabbit (cat # ab150077, Abcam) IgG secondary antibody (1:200 dilution) at room temperature for 1 hour. After rinsing and dehydration, slides were mounted using antifade mounting medium with DAPI (cat # H-1500-10, Vector Laboratories). An IgG secondary antibody only control was included for correction of background auto-immunofluorescence. The images were captured using a Zeiss LSM 800 with Airyscan microscope (Zeiss; Oberkochen, Germany). The fluorescence intensity was quantitated using ZEN-lite software (Zeiss) or Image J software (NIH). Fold change was calculated relative to vehicle-treated cells.

### Statistical analyses and reproducibility

All statistical analyses were conducted using GraphPad Prism 9.0 (La Jolla, CA) unless otherwise specified. Normal distribution of data was tested by the Shapiro-Wilk normality test. For normally distributed data, one-way ANOVA was used to test statistical significance for experiments with ≥ 3 groups followed by Tukey’s or Dunnett’s multiple comparison test. Student’s t-test was used for statistical significance comparison between two treatment groups. For data that failed to pass the normality test, the Kruskal-Wallis test was used to test statistical significance for experiments with ≥ 3 groups followed by Dunn’s post-hoc test. The nonparametric Mann-Whitney U test was used to examine statistical significance between two experimental conditions. All values are given as mean ± standard deviation (SD). Results are presented for experiments using at least three biological replicates, and *p* values < 0.05 were considered statistically significant. A detailed summary on statistical analyses is included in Source data.

### Reporting summary

Further information on research design is available in the Nature Portfolio Reporting Summary linked to this article.

### Data availability

All data that support the findings of this study are presented in this article and its Supplementary materials. All metabolomic profiling data were deposited at Metabolomics Workbench with accession IDs 4204, 4206, and 4207. Abbreviations and their corresponding full names are listed in alphabetical order in Supplementary Table 2. Reagents will be made available upon reasonable request to the corresponding author. Source data are provided with this paper.

## Acknowledgements

This work was supported in part by NIH grants R01 HL155107, R01 HL155096, R01 HL166137, and U01 HG007691; by AHA grant 957729; and by EU grant 101057619 to J.L; and by NIH grant R01 DK127278 to M.H. The authors acknowledge the Pulmonary Hypertension Breakthrough Initiative (PHBI) for providing human lung tissue, lung slides, PASMCs, and lung RNA samples, which is funded by NHLBI R24 HL123767 and the Cardiovascular Medical Research and Education Fund. We also thank Stephanie Tribuna for her expert technical assistance.

## Author contributions

W.X. and J.L. conceived, designed, and interpreted the study. W.X. performed experiments, analyzed and explained data. N.S. conducted experiments for the revision. W.M.O. and C.C. performed metabolomic profiling and data analysis. H.H. performed the L2HG measurement assay. S.W. and M.H. performed the PHD2 activity assay. B.M.W. and E.A. carried out immunohistochemistry fluorescence assay of rat lung tissue. W.X. and W.M.O. conducted the animal experiments. J.A.L. conducted the PAH cohort study and data analyses. M.H. and J.L. acquired funding support. J.L. supervised this project. All authors reviewed and approved this manuscript.

