## Supplemental figures and tables for "Branched chain α-ketoacids aerobically activate HIF1α signaling in vascular cells"

May 29, 2024

Divisions of Cardiovascular Medicine[1] and Pulmonary and Critical Care Medicine[1], Department of Medicine, Brigham and Women's Hospital and Harvard Medical School, Boston, MA 02115, USA; [2]Department of Toxicology, [3]Beijing Key Laboratory of Toxicological Research and Risk Assessment for Food Safety, and [4]Key Laboratory of State Administration of Traditional Chinese Medicine for Compatibility Toxicology, School of Public Health, Peking University, Beijing 100191, China; [5]Broad Institute of Massachusetts Institute of Technology and Harvard University, Cambridge, MA 02142, USA; [6]Department of Cell Biology, Blavatnik Institute, Harvard Medical School, Boston, MA 02115, USA

*Corresponding author:

Joseph Loscalzo, M.D., Ph.D.

Department of Medicine

Brigham and Women's Hospital

75 Francis St, Boston, MA 02115, USA

617-732-5127; 617-732-6439 (fax)

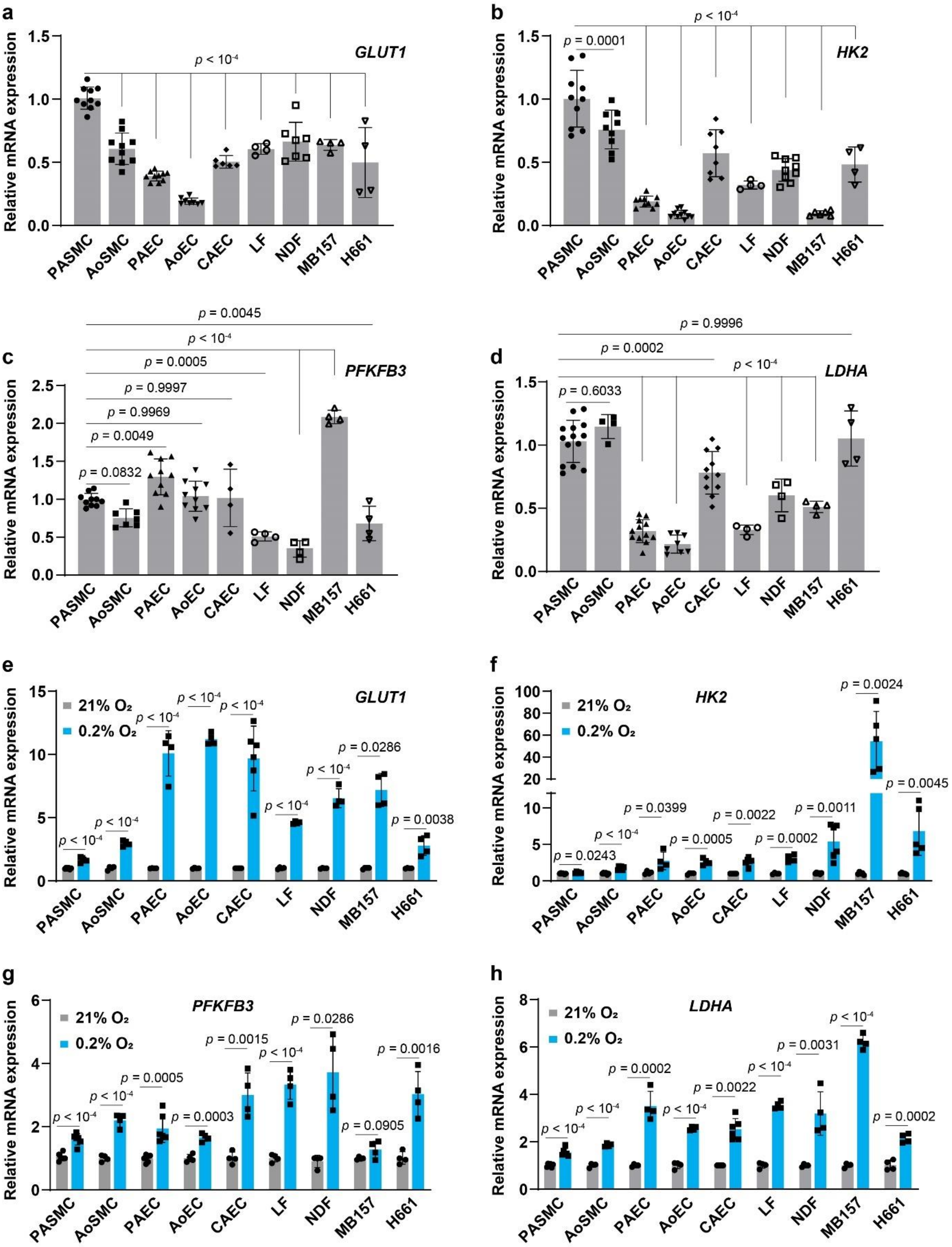
a
GLUT1
Relative mRNA expression
p < 10-4
PASMC AoSMC PAEC AoEC CAEC LF NDF MB157 H661
b
HK2
p < 10-4
p = 0.0001
c
PFKFB3
p = 0.0045
p < 10-4
p = 0.0005
p = 0.9997
p = 0.9969
p = 0.0049
p = 0.0832
d
LDHA
p = 0.9996
p = 0.0002
p < 10-4
p = 0.6033
e
GLUT1
21% O2
0.2% O2
p < 10-4
p = 0.0286
p = 0.0038
f
HK2
p = 0.0024
p = 0.0045
p = 0.0399
p = 0.0022
p = 0.0011
p = 0.0243
p = 0.0005
p = 0.0002
g
PFKFB3
p = 0.0286
p = 0.0015
p = 0.0016
p = 0.0005
p = 0.0003
p = 0.0905
h
LDHA
p = 0.0031
p = 0.0002
p = 0.0022

**Extended Data Fig.1 VSMCs exhibit high abundance of key glycolysis regulatory genes and are resistant to hypoxia-induced upregulation of these genes.**
**a-d,** Basal mRNA expression of glycolytic genes *GLUT1* (**a**), *HK2* (**b**), *PFKFB3* (**c**), and *LDHA* (**d**) in 9 different cell types under aerobic conditions. Fold change was calculated relative to PASMCs. *n* = 4-14.
**e-h,** mRNA expression of glycolytic genes in 9 different cell types cultured under normoxia (21% $O_2$) or hypoxia (0.2% $O_2$). Fold change was calculated relative to the corresponding type of cells grown under aerobic condition. *n* = 4-6.

One-way ANOVA followed by Dunnett's post-hoc test (**a-d**), Student's t test or Mann-Whitney U test (**e-h**) was applied when compared to untreated PASMCs (**a-d**) or normoxic cultures of the matched cell type (**e-h**).

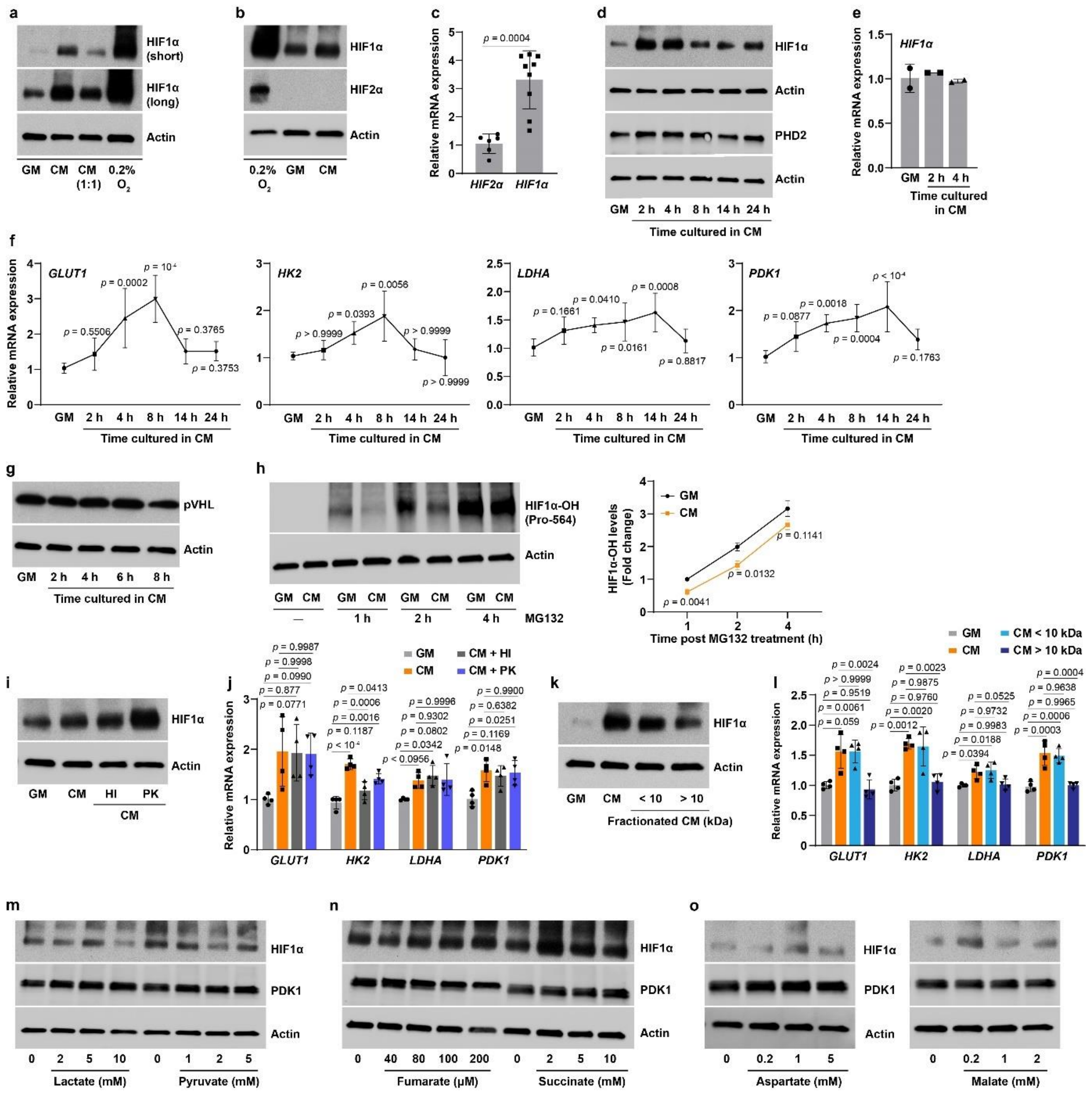

**Extended Data Fig. 2 Medium conditioned from PASMCs induces aerobic activation of HIF1α signaling.**

**a,** HIF1α protein expression in PASMCs cultured in growth medium (GM), conditioned medium (CM), or 1:1 (v/v) mix of CM and GM (CM 1:1). Cells grown under 0.2% $O_2$ were used as positive controls. Short (20 min) and long (60 min) represent film exposure duration.

**b,** HIF1α and HIF2α protein levels in cells cultured in GM, CM, or 0.2% $O_2$.

**c,** *HIF1α* and *HIF2α* mRNA abundance in PASMCs cultured in GM. Fold change was calculated relative to HIF2α. $n$ = 6-9.

**d**, HIF1α and PHD2 protein levels in PASMCs cultured in CM for 2-24 hours.

**e,** *HIF1α* mRNA expression in cells cultured in CM for 2 and 4 hours. Fold change was calculated relative to PASMCs grown in GM. $n$ = 2.

**f,** mRNA expression of HIF1α target genes in glucose metabolism in PASMCs when cultured in CM for various time points. Fold change was calculated relative to cells in GM. *n* = 6.
**g,** von-Hippel Lindau protein (pVHL) levels in cells cultured in CM for different times.
**h,** Representative immunoblots and quantitation of hydroxylated HIF1α protein (HIF1α-OH Pro-564) levels in GM- or CM-cultured PASMCs after addition of proteasomal inhibitor MG132 (20 μM) for 1-4 hours. Fold change was calculated relative to GM-cultured PASMCs at 1 hour of MG132 incubation. *n* = 3.
**i,j** HIF1α protein levels (**i**) and mRNA expression of its transcriptional targets (**j**) in GM, CM, and heat inactivation (HI) or proteinase K (PK) treated CM cultured PASMCs. Fold change in **j** was calculated relative to GM-cultured cells. *n* = 4 (**j**).
**k,l** HIF1α protein levels (**k**) and mRNA expression of its transcriptional targets (**l**) in GM, CM, and fractionated CM (larger than 10 kDa fraction, >10 and less than 10 kDa fraction, < 10) cultured PASMCs. Fold change in **l** was calculated relative to GM-cultured cells. *n* = 4 (**l**).
**m-o,** HIF1α and its target PDK1 protein levels in PASMCs treated with lactate or pyruvate (**m**), fumarate or succinate (**n**), aspartate or malate (**o**) as indicated doses.

Mann-Whitney U test (**c**), Student t test (**h**), or one-way ANOVA followed by Tukey's post-hoc test or Kruskal-Wallis test followed by Dunn's test (**f**, **j**, **l**) was applied when compared to HIF2α in PASMCs (**c**), to GM-cultured PASMCs with time-matched MG132 treatment (**h**) or no treatment (**f**, **j**, **l**) or to CM-cultured PASMCs (**j**, **l**).

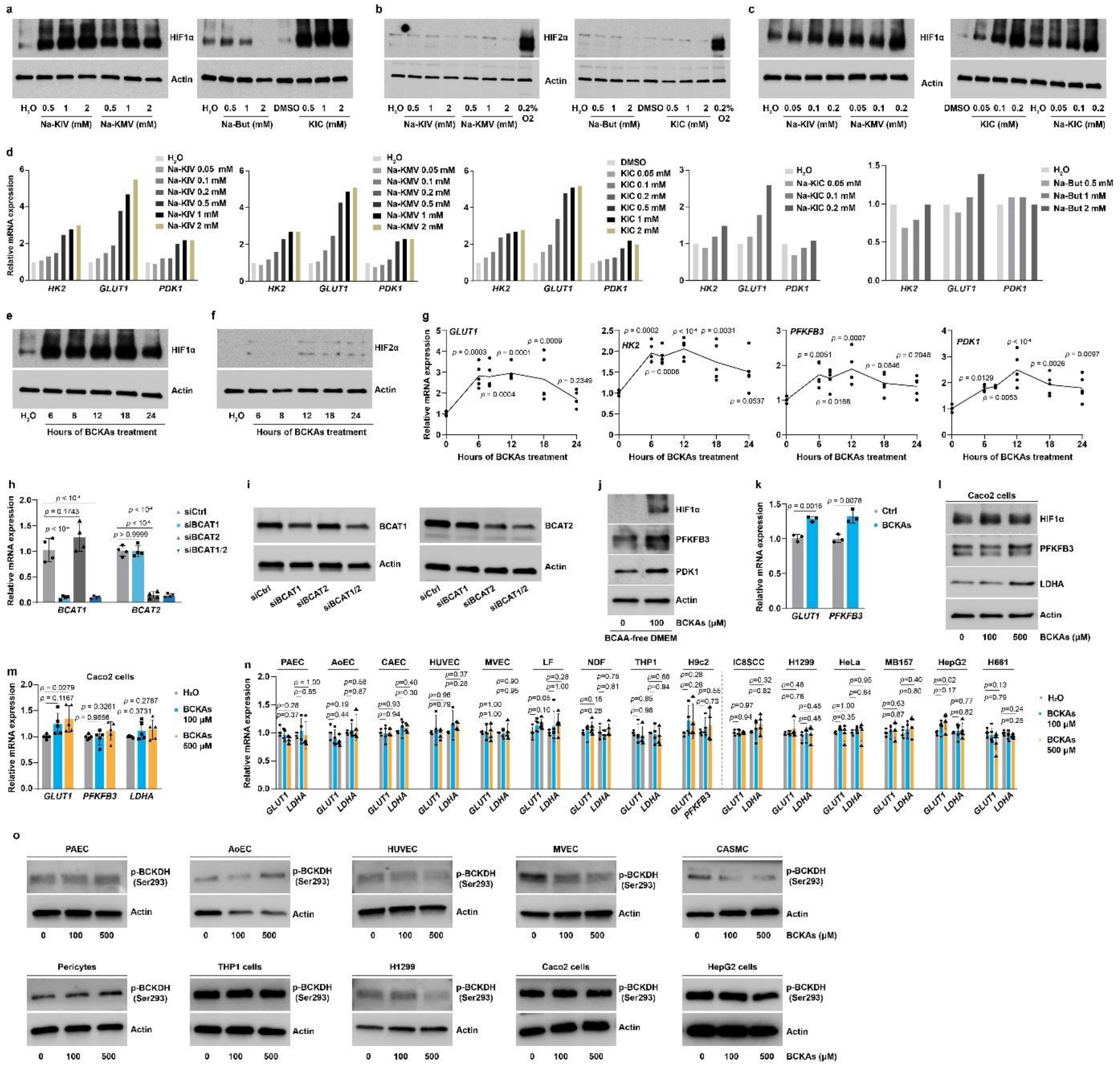

**Extended Data Fig.3 BCKAs are the mediators of paracrine activation of HIF1α signaling under aerobic conditions.**

**a-c,** HIF1α (**a, c**) and HIF2α (**b**) proteins in PASMCs treated with 0.05-2 mM of sodium salts of KIV (Na-KIV), KMV (Na-KMV), butyrate (Na-But), KIC (Na-KIC), or acid form of KIC (KIC; DMSO as vehicle control) for 8 hours. Hypoxia (0.2% $O_2$) induced HIF2α protein stabilization was included for comparison.

**d,** mRNA expression of three HIF1α target genes in glucose metabolism in PASMCs stimulated with 0.05-2 mM of Na-KIV, Na-KMV, Na-KIC, KIC, and Na-But for 8 hours. Fold change was calculated relative to vehicle control ($H_2O$ or DMSO) treated cells. *n* = 1.

**e-g,** HIF1α (**e**) and HIF2α (**f**) protein levels and the mRNA expression of HIF1α target genes in glucose metabolism (**g**) in PASMCs stimulated with BCKAs (100 µM Na-KIC, 50 µM of each Na-KIV and Na-KMV) for different time points. Fold change in **g** was calculated relative to untreated control cells at 8-hour time point. *n* = 5 (**g**).

**h,i,** mRNA (**h**) and protein (**i**) expression of BCAT1 and BCAT2 in PASMCs transfected with siRNAs for control (siCtrl), *BCAT1* (siBCAT1), *BCAT2* (siBCAT2), or both (siBCAT1/2). Fold change in **h** was relative to siCtrl-transfected cells. *n* = 4 (**h**).
**j,k,** Protein levels of HIF1α, PFKFB3, and PDK1 (**j**) and mRNA expression of *GLUT1* and *PFKFB3* (**k**) in PASMCs cultured in BCAA-free DMEM in the presence or absence of BCKAs for 8 hours. Fold change in **k** was calculated relative to untreated control cells. *n* = 3 (**k**).
**l,m,** HIF1α, PFKFB3, and LDHA protein levels (**l**) and the mRNA expression of HIF1α regulatory genes in glycolysis (**m**) of human colorectal adenocarcinoma Caco2 cells treated with BCKAs. Fold change in (**m**) was relative to vehicle control treated cells. *n* = 5 (**m**).
**n,** mRNA expression of *GLUT1*, *LDHA*, or *PFKFB3* in normal and cancerous cells after stimulation with different doses of BCKAs. Fold change was calculated relative to their own untreated control cells. Dotted line separates normal *vs.* malignant cell types. *n* = 4-5.
**o**, Phosphorylated BCKDH (p-BCKDH) protein levels in 10 different types of cells with BCKA treatment.

One-way ANOVA followed by Dunnett's post-hoc test (**g**, **h**, **m**, **n**) or Student's t test (**k**) was applied when compared to untreated control PASMCs (**g**), siCtrl-transfected PASMCs (**h**), BCAA-free DMEM-cultured control PASMCs (**k**), untreated Caco2 cells (**m**), or the corresponding untreated cells (**n**).

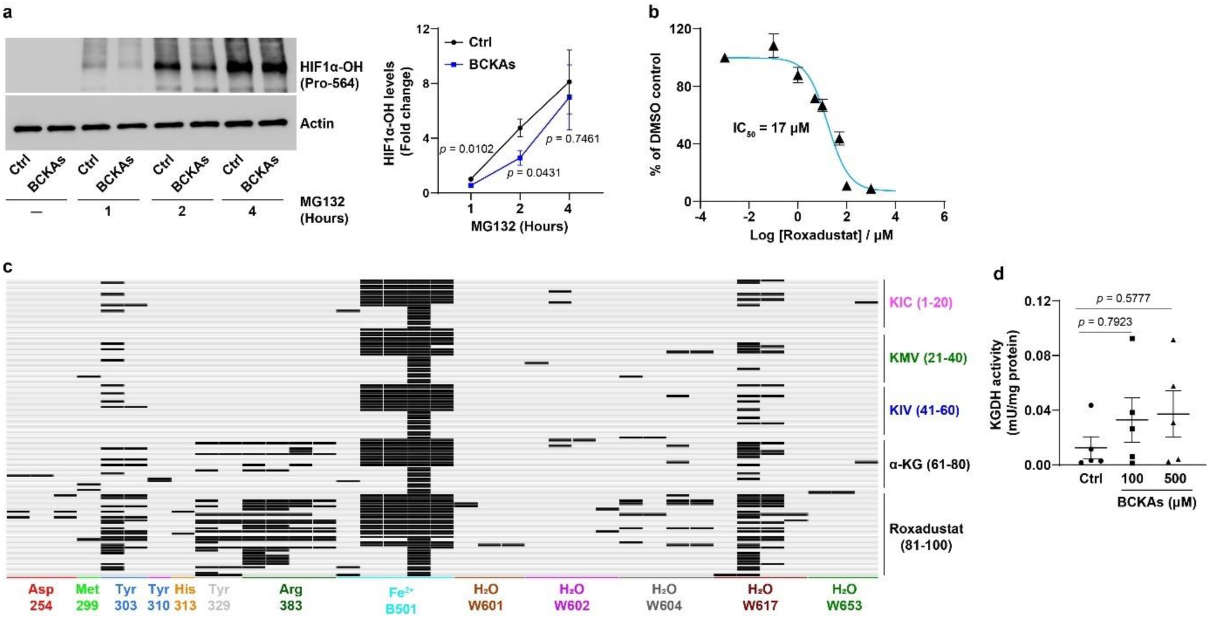


**Extended Data Fig. 4 Effects of BCKAs on PHD2 and KGDH activity.**

**a,** PASMCs were treated with BCKAs (100 µM of KIC, 50 µM of each KIV and KMV) or vehicle control for 8 hours followed by addition of proteasomal inhibitor MG132 (20 µM) for 1-4 hours. Hydroxylated HIF1α (HIF1α-OH Pro-564) protein levels were measured and quantitated. Fold change was calculated relative to untreated cells with MG132 incubation for 1 hour. $n = 3$.

**b,** Inhibition curve and $IC_{50}$ value of roxadustat for PHD2 hydroxylase activity. $n = 4$.

**c,** Protein-ligand interaction fingerprint (PLIF) prediction of 20 potential binding configurations of each BCKA with the PHD2 active site. α-KG and roxadustat, two known ligands of PHD2 enzyme, were included for comparison.

**d,** KGDH activity in PASMCs treated with BCKAs. $n = 5$.

Student's t test (**a**) or Kruskal-Wallis test followed by Dunn's post-hoc test (**d**) was applied when compared to untreated PASMCs at time-matched MG132 treatment (**a**) or untreated PASMCs (**d**).

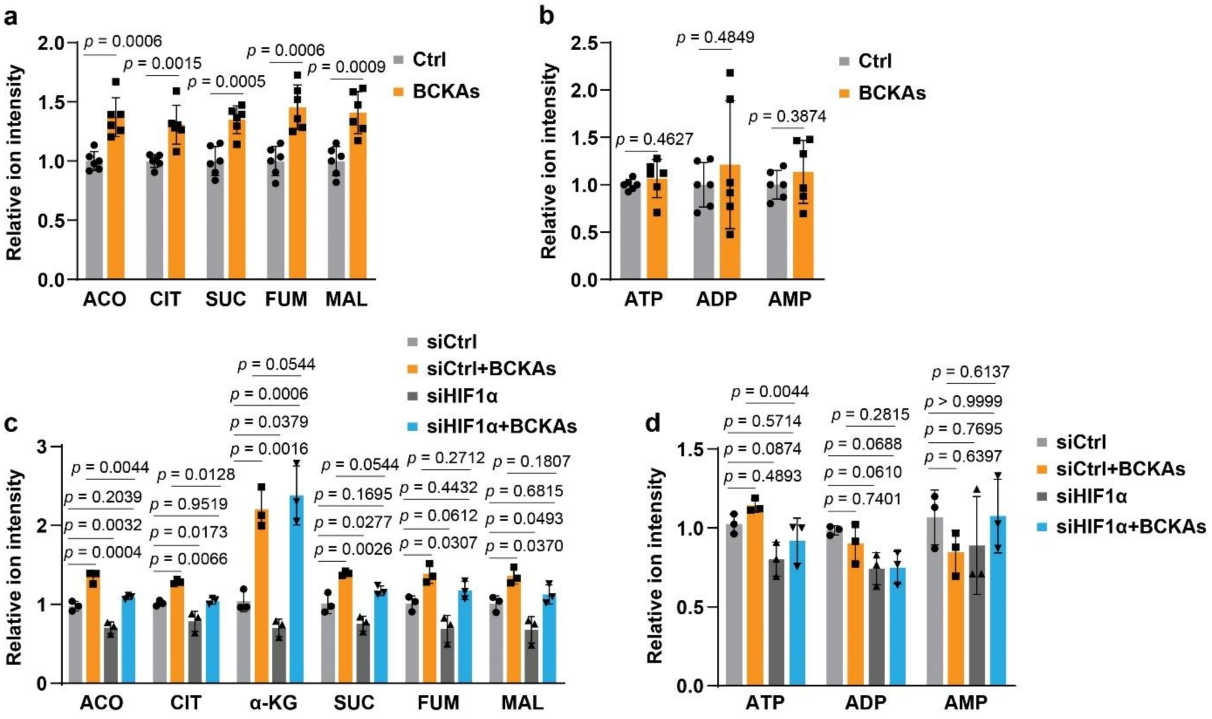


**Extended Data Fig. 5 The influence of BCKAs on mitochondrial respiration and its dependence on HIF1α activity in PASMCs.**

**a,b,** LC-MS measurements of intermediary metabolites aconitate (ACO), citrate (CIT), succinate (SUC), fumarate (FUM), and malate (MAL) of the TCA cycle (**a**), and of ATP and its derivatives (**b**) in PASMCs in the presence or absence of BCKAs. Fold change was calculated relative to control cells. $n = 6$.

**c,d,** PASMCs were transfected with human *HIF1α* siRNA (siHIF1α) or control siRNA (siCtrl) followed by treatment with BCKAs. LC-MS was used to measure the TCA cycle metabolites (**c**), and ATP and its metabolites (**d**). Fold change was calculated relative to siCtrl-transfected and untreated cells. $n = 3$.

Student's t test (**a**, **b**) or one-way ANOVA followed by Tukey's post-hoc test (**c**, **d**) was applied when compared to control PASMCs (**a**, **b**), or siCtrl-transfected and control or BCKA-treated cells (**c**, **d**).

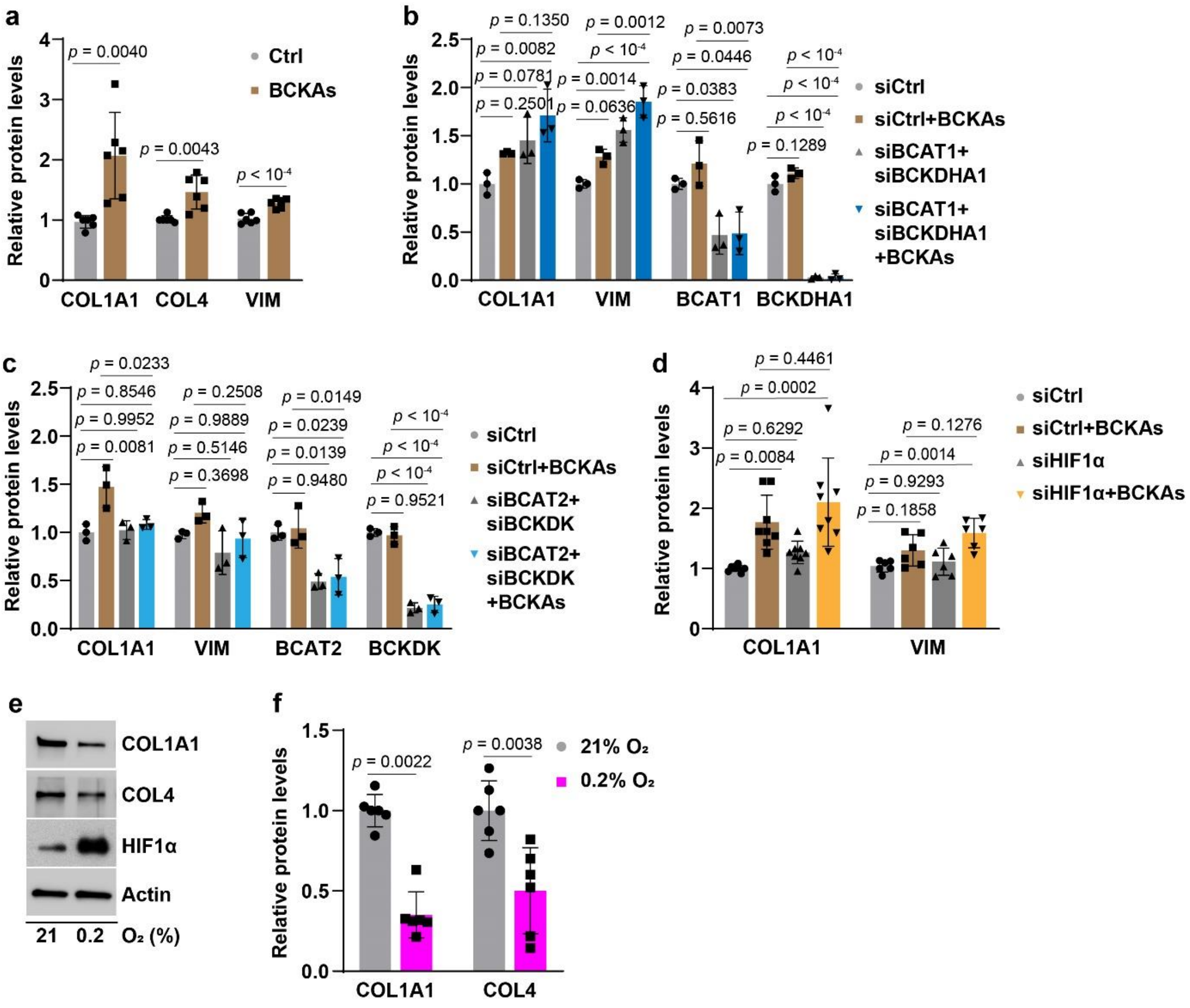


**Extended Data Fig. 6 The levels of synthetic phenotype marker proteins in PASMCs.**

**(a)** Protein levels in PASMCs treated with BCKAs. Fold change was calculated relative to untreated control. *n* = 6.

**(b-d)** Protein levels in PASMCs transfected with control siRNA (siCtrl), *BCAT1* and *BCKDHA1* siRNA (siBCAT1+siBCKDHA1; **b**), or *BCAT2* and *BCKDK* siRNA (siBCAT2+siBCKDK; **c**), or *HIF1α* siRNA (siHIF1α; d) with or without BCKA treatment. Fold change was calculated relative to siCtrl-transfected and untreated control. *n* = 3 (**b, c**) and 8 (**d**).

(**e**,**f**) Representative immunoblots (**e**) and quantitation (**f**) of COL1A1 and COL4 protein levels in PASMCs cultured in 21% $O_2$ or 0.2% $O_2$. HIF1α protein was included as a positive control in hypoxia. Fold change in **f** was relative to normoxic cultures of PASMCs. *n* = 6.

Student's t test (COL1A1 and VIM in **a**, COL4 in **f**), Mann-Whitney U test (COL4 in **a**, COL1A1 in **f**), or one-way ANOVA followed by Tukey's post-hoc test (**b-d**) was applied when compared to control PASMCs (**a**, **f**), or siCtrl-transfected and control or BCKA-treated cells (**b-d**).

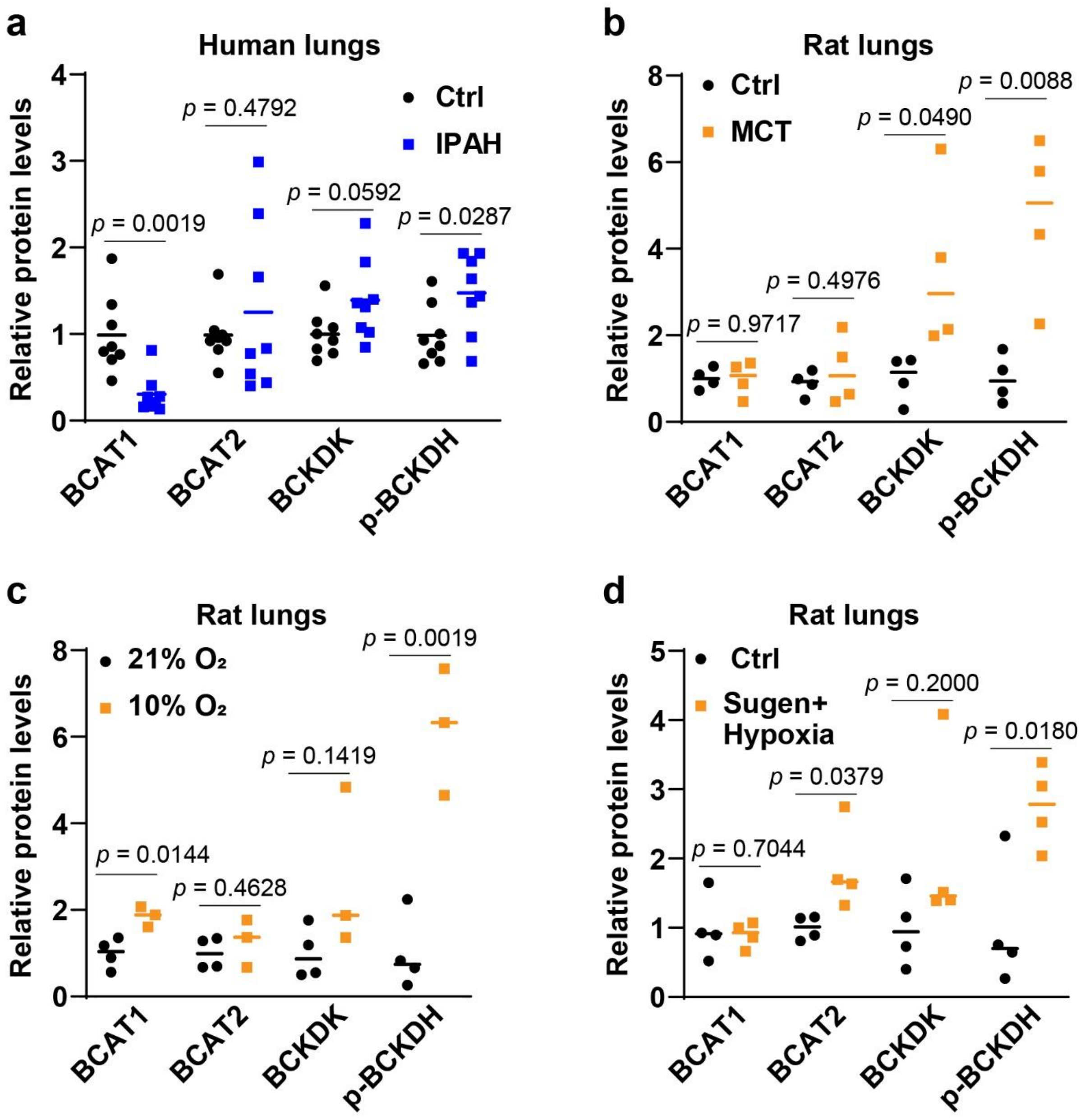


**Extended Data Fig. 7 The levels of key BCKA metabolic proteins in the lungs of PAH patients and rats.**

**a,** Quantitation results of BCAT1, BCAT2, BCKDK, and p-BCKDH proteins in the lungs of IPAH patients. *n* = 8 individuals.

**b-d,** Quantitation results of BCAT1, BCAT2, BCKDK, and p-BCKDH proteins in the lungs of PAH rats treated with MCT (**b**), hypoxia (10% $O_2$; **c**), or Sugen5416+hypoxia (**d**). *n* = 3-4 rats.

Student's t test (**a-d**) or Mann-Whitney U test (BCAT1 in **a** and BCKDK in **d**) was applied when compared to control patients (**a**) or animals (**c-d**).

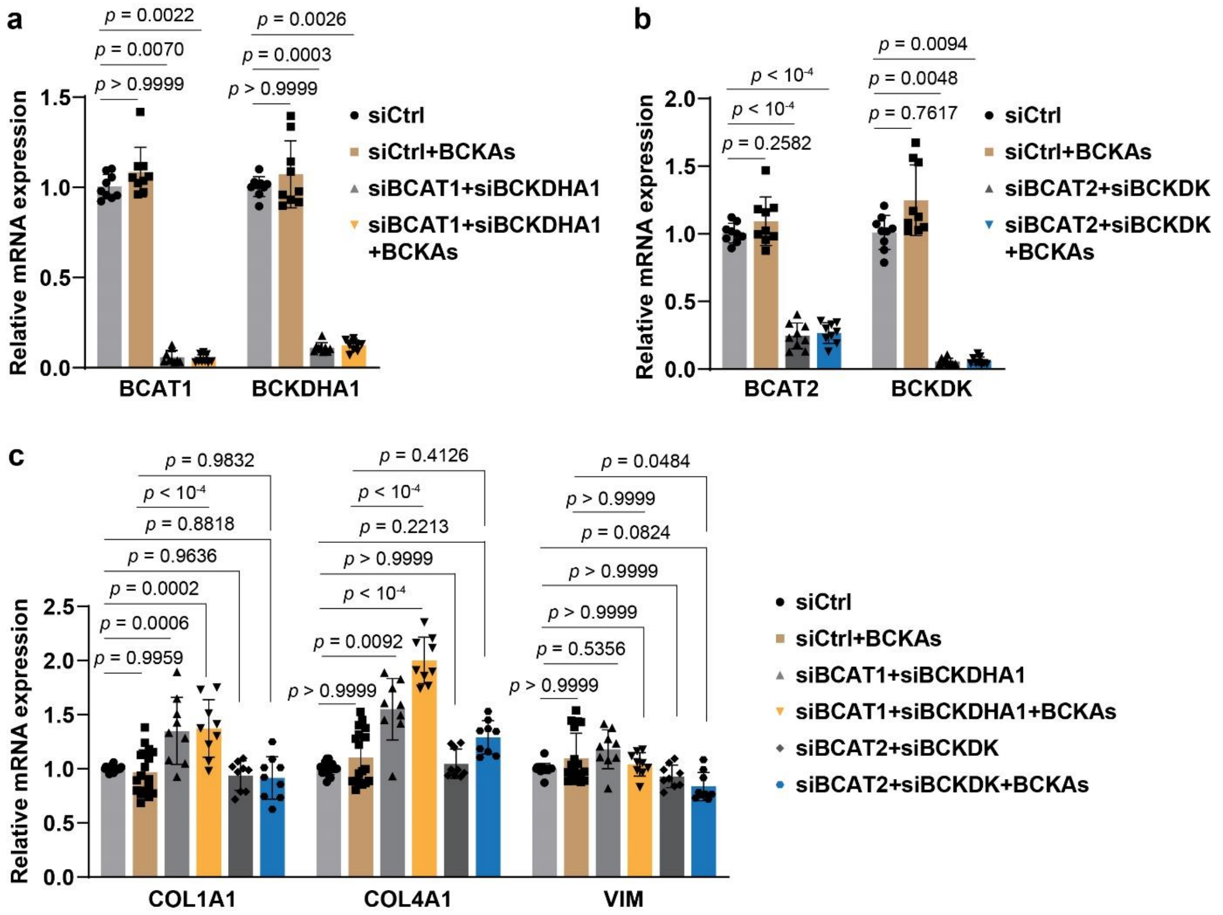


**Extended Data Fig. 8 The expression of synthetic marker genes in IPAH-PASMCs with endogenous and exogenous manipulation of BCKA levels.**

(**a**) mRNA expression of BCAT1 and BCKDHA1 in IPAH-PASMCs transfected with siRNAs for control (siCtrl) or *BCAT1* and *BCKDHA1* (siBCAT1+siBCKDHA1) followed by BCKAs or vehicle control treatment. $n = 9$ from 3 individuals.

(**b**) mRNA expression of BCAT2 and BCKDK in IPAH-PASMCs transfected with siCtrl or *BCAT2* and *BCKDK* (siBCAT2+siBCKDK) followed by BCKAs or vehicle control treatment. $n = 9$ from 3 individuals.

(**c**) mRNA expression of synthetic marker genes in IPAH-PASMCs transfected and treated as described in panels **a** and **b**. $n$ = 9-18 from 3 individuals.

Kruskal-Wallis test followed by Dunn's post-hoc test (**a-c**) or one-way ANOVA followed by Dunnett's (**b**) or Tukey's (**c**) post-hoc test was applied when compared to siCtrl-transfected control cells (**a-c**) or BCKA-treated cells (**c**).

**Supplementary Table 1 Clinical and demographic information on human specimen presented in this study**

| Patient ID | Clinical diagnosis | Gender | Race | Ethnicity |
|---|---|---|---|---|
| **Lung RNA samples** | | | | |
| AH-007 | FDL | M | White | Non-Hispanic |
| AH-009 | FDL | M | White | Non-Hispanic |
| AH-012 | FDL | M | White | Non-Hispanic |
| AH-013 | FDL | F | White | Non-Hispanic |
| BA-033 | FDL | M | White | Non-Hispanic |
| BA-040 | FDL | M | Unknown | Hispanic or Latino |
| BA-046 | FDL | F | Unknown | Hispanic or Latino |
| BA-055 | FDL | M | White | Non-Hispanic |
| UC-010 | FDL | F | White | Non-Hispanic |
| VA-005 | FDL | M | White | Non-Hispanic |
| BA-017 | IPAH | F | White | Non-Hispanic |
| CC-017 | IPAH | M | White | Non-Hispanic |
| CC-030 | IPAH | F | White | Non-Hispanic |
| ST-004 | IPAH | F | White | Non-Hispanic |
| ST-010 | IPAH | M | White | Non-Hispanic |
| ST-017 | IPAH | M | White | Non-Hispanic |
| ST-019 | IPAH | M | White | Hispanic or Latino |
| ST-042 | IPAH | M | White | Non-Hispanic |
| UA-013 | IPAH | M | Asian | Non-Hispanic |
| VA-015 | IPAH | F | White | Non-Hispanic |
| **Frozen lung tissues** | | | | |
| AH-012 | FDL | M | White | Non-Hispanic |
| AH-013 | FDL | F | White | Non-Hispanic |
| AH-016 | FDL | M | White | Non-Hispanic |
| BA-040 | FDL | M | Unknown | Hispanic or Latino |
| BA-043 | FDL | M | Unknown | Hispanic or Latino |
| BA-046 | FDL | F | Unknown | Hispanic or Latino |
| BA-048 | FDL | M | White | Non-Hispanic |
| BA-049 | FDL | M | Unknown | Non-Hispanic |

| | | | | |
|---|---|---|---|---|
| BA-055 | FDL | M | White | Non-Hispanic |
| BA-062 | FDL | M | Asian | Unknown |
| BA-017 | IPAH | F | White | Non-Hispanic |
| CC-030 | IPAH | F | White | Non-Hispanic |
| ST-028 | IPAH | F | White | Hispanic or Latino |
| ST-033 | IPAH | F | White | Non-Hispanic |
| ST-037 | IPAH | F | Unknown | Hispanic or Latino |
| ST-042 | IPAH | M | White | Non-Hispanic |
| ST-052 | IPAH | M | Asian | Non-Hispanic |
| UA-013 | IPAH | M | Asian | Non-Hispanic |
| VA-011 | IPAH | F | White | Non-Hispanic |
| VA-015 | IPAH | F | White | Non-Hispanic |
| **Lung slides** | | | | |
| BA-049 | FDL | M | Unknown | Non-Hispanic |
| BA-062 | FDL | M | Asian | Unknown |
| BA-046 | FDL | F | Unknown | Hispanic or Latino |

**PASMCs**

| Patient ID | Clinical diagnosis | Gender | Race | Age (Y) |
|---|---|---|---|---|
| CC-013 | IPAH | F | White | 27 |
| ST-019 | IPAH | M | White | 25 |
| ST-026 | IPAH | M | White | 40 |
| UA-013 | IPAH | M | Asian | 18 |
| VA-011 | IPAH | F | White | 32 |

FDL: failed donor lung; IPAH: Idiopathic pulmonary arterial hypertension

**Supplementary Table 2 Abbreviations and their corresponding full names used**

| Abbreviation | Full name |
|---|---|
| α-KG | α-ketoglutarate |
| ACTA2 | α-smooth muscle actin |
| AoSMCs | aortic smooth muscle cells |
| BCAAs | branched chain amino acids |
| BCAT | branched chain amino acid transaminase |
| BCKAs | branched chain α-ketoacids |
| BCKDH | branched chain ketoacid dehydrogenase complex |
| BCKDK | branched chain ketoacid dehydrogenase kinase |
| CASMCs | coronary artery smooth muscle cells |
| COL1A1 | collagen 1A1 |
| COL4 | collagen 4 |
| ECAR | extracellular acidification rate |
| GLUT1 | glucose transporter 1 |
| HK2 | hexokinase 2 |
| HIF1α | hypoxia-inducible factor 1α |
| KGDH | α-KG dehydrogenase |
| KIC | α-ketoisocaproate |
| KIV | α-ketoisovalerate |
| KMV | α-keto-β-methylvalerate |
| L2HG | L-2-hydroxyglutarate |
| L2HGDH | L2HG dehydrogenase |
| LDHA | lactate dehydrogenase A |
| mPAP | mean pulmonary artery pressure |
| OCR | oxygen consumption rate |
| PAH | pulmonary arterial hypertension |
| PASMCs | pulmonary arterial smooth muscle cells |
| PDK1 | pyruvate dehydrogenase kinase 1 |
| PFKFB3 | 6-phosphofructo-2-kinase/fructose 2,6-biphosphatase 3 |
| PHD2 | prolyl hydroxylase domain-containing protein 2 |
| PVR | pulmonary vascular resistance |
| ROS | reactive oxygen species |
| TCA | tricarboxylic acid |
| VIM | vimentin |
| VSMCs | vascular smooth muscle cells |
